# Elements at the 5′ end of *Xist* harbor SPEN-independent transcriptional antiterminator activity

**DOI:** 10.1101/2020.05.13.090506

**Authors:** Jackson B. Trotman, David M. Lee, Rachel E. Cherney, Sue O. Kim, Kaoru Inoue, Megan D. Schertzer, Steven R. Bischoff, Dale O. Cowley, J. Mauro Calabrese

**Author notes:** These authors contributed equally to this work (Jackson B. Trotman, David M. Lee).

## Abstract

The *Xist* lncRNA requires Repeat A, a conserved RNA element located in its 5′ end, to induce gene silencing during X-chromosome inactivation. Intriguingly, Repeat A is also required for the production of *Xist*. While silencing by Repeat A requires the protein SPEN, how Repeat A promotes *Xist* production remains unclear. We report that in mouse embryonic stem cells, expression of a transgene comprising the first two kilobases of *Xist* (*Xist*-2kb) causes transcriptional readthrough of multiple downstream polyadenylation sequences. Readthrough required Repeat A and the ~750 nucleotides downstream but did not require SPEN. Despite associating with SPEN and chromatin, *Xist*-2kb did not robustly silence transcription, whereas a transgene comprising *Xist*’s first 5.5 kilobases robustly silenced transcription and read through its polyadenylation sequence. Longer, spliced *Xist* transgenes also induced robust silencing yet terminated efficiently. Thus, in contexts examined here, *Xist* requires sequence elements beyond its first two kilobases to robustly silence transcription, and the 5′ end of *Xist* harbors SPEN-independent transcriptional antiterminator activity that can repress proximal cleavage and polyadenylation. In endogenous contexts, this antiterminator activity may help produce full-length *Xist* RNA while rendering the *Xist* locus resistant to silencing by the same repressive complexes that the lncRNA recruits to other genes.

## Introduction

The long noncoding RNA (lncRNA) *Xist* functions *in cis* to silence nearly all genes along the 165-megabase (Mb) X chromosome as part of X-chromosome inactivation (XCI), the dosage compensation process that occurs early during the development of eutherian mammals. At the level of sequence composition, *Xist* is notable for the presence of several internal domains of tandem repeats, which each help *Xist* achieve its repressive function by recruiting distinct subsets of RNA-binding proteins (1–22). One such repeat is found in the first thousand nucleotides of *Xist* and is called “Repeat A”. Repeat A consists of eight to nine tandemly arrayed, 50-nucleotide-long repeating elements that each harbor a degenerate U-rich region followed by a distinctive GC-rich region that is highly conserved among eutherians (1,2,4,8).

Gene silencing induced by *Xist* has been shown to depend on Repeat A as well as the Repeat A-binding protein SPEN. Recruitment of SPEN to the X chromosome via Repeat A likely induces gene silencing by locally activating and/or recruiting various corepressor and histone deacetylase complexes (10,23–25). Concordantly, deletion of Repeat A or SPEN each results in failure of XCI (4,7,10,23–29), and robust associations between SPEN and Repeat A have been detected *in vivo* (10,21,30). *In vitro*, SPEN binds to single-stranded regions of Repeat A that are located directly adjacent to its structured, GC-rich segments (13,28).

Although Repeat A and SPEN are both necessary for XCI, *Xist* transgenes that contain Repeat A but lack essentially all other *Xist* sequence downstream are unable to induce chromosome-level silencing, implying either (i) that SPEN requires sequence elements in addition to Repeat A to associate with *Xist* or (ii) that *Xist* sequence elements in addition to a SPEN-bound Repeat A are needed to induce chromosome-level silencing (4,8,16).

Moreover, despite clear links between SPEN and Repeat A, questions remain regarding the mechanism through which Repeat A carries out its functions. Most notably, deletion of Repeat A from the endogenous *Xist* locus causes not only a failure of XCI but a dramatic reduction in the abundance of full-length, spliced *Xist* (7,20,29,31). Similar reductions in *Xist* RNA abundance have been observed upon Repeat A deletion in transgenic contexts (27,32–34), seemingly due to a defect in the production of nascent *Xist* and not due to a reduction in RNA stability (33). Current evidence is at odds as to whether the loss of SPEN reduces or has no impact on *Xist* RNA abundance (25,27). Similarly, a seminal study found that the repressive function of an *Xist* transgene lacking Repeat A can be restored by appending Repeat A to its 3′ end, suggesting that the function of Repeat A is independent of its position within *Xist* (4). In contrast, a recent report found loss of *Xist* abundance and *Xist*-induced silencing when Repeat A was moved from the 5′ end of *Xist* to positions further downstream, suggesting the opposite conclusion – that in addition to the sequence of Repeat A, its 5′-proximal position may be important for the production and function of *Xist* (35). The basis for this apparent difference is unclear.

We recently developed a transgenic assay that recapitulates certain aspects of Repeat A-dependent gene silencing, which we called TETRIS (<u>T</u>ransposable <u>E</u>lement to <u>T</u>est <u>R</u>NA’s effect on transcription *in c<u>is</u>*; (36)). In TETRIS, a doxycycline-inducible transgene that comprises the first two kilobases (kb) of *Xist* and contains Repeat A (*Xist*-2kb) is positioned in convergent orientation relative to a luciferase reporter gene in the context of a piggyBac cargo plasmid. After insertion of the plasmid into chromatin via the piggyBac transposase, induction of *Xist*-2kb expression results in an 80 to 90% reduction of luciferase activity relative to uninduced cells. We demonstrated that repression of luciferase activity by *Xist*-2kb in this assay requires the same sequence motifs within Repeat A that are required for transcriptional repression by full-length *Xist* – its GC-rich portions but not its U-rich spacer sequences – as well an adjacent region that harbors structured elements and intervening sequence (4,14,36).

Given the unresolved questions surrounding Repeat A, we sought to investigate the mechanism of repression induced by *Xist*-2kb in the TETRIS assay as well as in a transgenic, single-copy insertion assay that is analogous to assays previously employed to identify seminal aspects of *Xist* biology (4,8,10,14,16,19,23–28,32). Through a circuitous series of experiments, we unexpectedly found that in these transgenic contexts, Repeat A and nearby sequence harbor transcriptional antiterminator activity that does not require the silencing cofactor SPEN. Furthermore, despite associating with SPEN and with chromatin, RNA produced from the *Xist*-2kb transgene was unable to induce robust local or long-distance transcriptional silencing. Instead, long-distance silencing required synergy between elements located in the first 2kb of *Xist* and regions downstream. Our findings have implications for understanding how both long-distance silencing and suppression of premature polyadenylation are carried out by *Xist* and possibly other RNAs.

## Materials and Methods

### Embryonic stem cell culture

E14 mouse embryonic stem cells (ESCs; kind gift of D. Ciavatta) were cultured using standard methods. Complete details can be found in the Supplementary Methods.

### TETRIS line generation

TETRIS lines were made as described in (36). Briefly, 500,000 E14 cells were seeded in a single well of a 6-well plate and transfected 24 h later with 0.5 μg TETRIS cargo plasmid, 0.5 μg *rtTA*-cargo plasmid, and 1 μg of pUC19-piggyBAC transposase plasmid using Lipofectamine 3000 (Invitrogen) according to manufacturer instructions. Cells were selected for 7-9 days with puromycin (2 μg/mL) and G418 (200 μg/mL) beginning 24 h after transfection.

### TETRIS luminescence assays

For each independent TETRIS cell line, six wells of a 24-well plate were seeded at 100,000 cells per well. Three of the six wells were induced with 1 μg/mL doxycycline (Sigma) beginning when the cells were plated; the remaining three wells served as “no dox” controls. After 48 h, the cells were washed with PBS and lysed with 100 μL of passive lysis buffer (Promega) and luciferase activity was measured using Bright-Glo Luciferase Assay reagents (Promega) on a PHERAstar FS plate reader (BMG Labtech). Luciferase activity was normalized to total protein concentration in the lysates measured via Bradford assay (Bio-Rad). See Supplementary Table S1 for additional information regarding experimental replicates.

### RNA isolation, subcellular fractionation, and RT-qPCR

RNA was isolated using Trizol according to manufacturer protocol (Invitrogen). Subcellular fractionation was performed essentially as in (37,38). For RT-qPCR assays, equal amounts of RNA (0.5-1 μg) were reverse transcribed using the High-Capacity cDNA Reverse Transcription Kit (Applied Biosystems) with a pool of random primers or a single, strand-specific primer (Supplementary Table S2). qPCR was performed using iTaq Universal SYBR Green (Bio-Rad) and custom primers (Supplementary Table S2) on a Bio-Rad CFX96 system with the following thermocycling parameters: initial denaturation at 95 °C for 10 min; 40 cycles of 95 °C for 15 s, 60 °C for 30 s, and 72 °C for 30 s followed by a plate read. Data were normalized (usually to the average of the “no dox” control measurements) using Microsoft Excel and plotted using GraphPad Prism 8. See Supplementary Table S1 for information regarding experimental replicates. Complete details can be found in the Supplementary Methods.

### Stellaris single-molecule sensitivity RNA FISH

Custom Stellaris FISH probes were designed against the first 2kb of *Xist*, firefly luciferase (*luc2* in pGL4.10; Promega), and the hygromycin resistance gene (*HygroR*) using the Stellaris RNA FISH Probe Designer (Biosearch Technologies, Inc.) and labeled with Quasar 670 (*Xist*-2kb) or 570 (*Luc* and *HygroR*) dye. FISH was performed as described in (39). Complete details can be found in the Supplementary Methods.

### Northern blots

Northern blots were performed using a protocol adapted from (40). Complete details can be found in the Supplementary Methods.

### Recombinase-mediated cassette exchange (RMCE)

Complete details can be found in the Supplementary Methods. Briefly, a male F1-hybrid mouse ESC line (derived from a cross between C57BL/6J (B6) and CAST/EiJ (Cast) mice; kind gift of T. Magnuson) was made competent for RMCE by insertion of a custom homing cassette into the *Rosa26* locus on chromosome 6 via homologous recombination. *Xist* transgenes were cloned via PCR or recombineering into a custom RMCE-cargo vector and then electroporated along with a plasmid expressing Cre-recombinase into RMCE-competent cells using a Neon Transfection System (Invitrogen). Individual colonies were picked and genotyped, then rendered doxycycline-sensitive using the *rtTA*-expression cassette from (41).

### RNA sequencing and analysis

RNA-seq libraries were prepared using the RNA HyperPrep Kit with RiboErase (Kapa Biosciences) and sequenced on an Illumina NextSeq 500 machine using a 75-cycle high output NextSeq kit (Illumina). Sequencing reads were aligned and processed essentially as in (42,43). Differential expression analysis was performed with DESeq2 (44). Complete details can be found in the Supplementary Methods.

### RNA immunoprecipitation

RNA immunoprecipitation experiments were performed using a protocol from (45). Complete details can be found in the Supplementary Methods.

## Results

### *Xist*-2kb represses expression of adjacent genes by reading through multiple polyadenylation sequences

TETRIS is a transgenic assay that allows the sequence of a lncRNA to be manipulated in a plasmid and then tested for its ability to repress adjacent reporter genes in a chromosomal context. The assay employs the piggyBac transposase to insert a cassette containing a doxycycline-inducible lncRNA gene, a luciferase reporter gene (*Luc*), and a puromycin resistance gene (*PuroR*) into the genomes of transfected cells (Figure 1A). Expression of non-repressive lncRNAs in TETRIS, such as *Hottip* (Figure 1B), typically causes a ~1.5-fold increase in luciferase activity, a mild enhancer effect that we attribute to the proximity of the doxycycline-inducible TRE promoter and the *PGK* promoter that drives *Luc* gene expression (36). In contrast, induced expression of the first 2 kb of *Xist* (*Xist*-2kb) in TETRIS causes an 80 to 90% reduction of luciferase activity, which we confirmed by observing a loss of luciferase protein by western blot (Figure 1B). We also found that *Xist*-2kb induction repressed *PuroR* expression, impairing cell survival in the presence of puromycin (Figure 1C). Previously, we found that silencing by *Xist*-2kb depends on Repeat A and an additional ~750 nucleotides of sequence located just downstream, implying that the mechanisms of *Xist*-2kb-induced repression in TETRIS are related to those employed by *Xist* in endogenous contexts (4,36).

**Figure 1.**
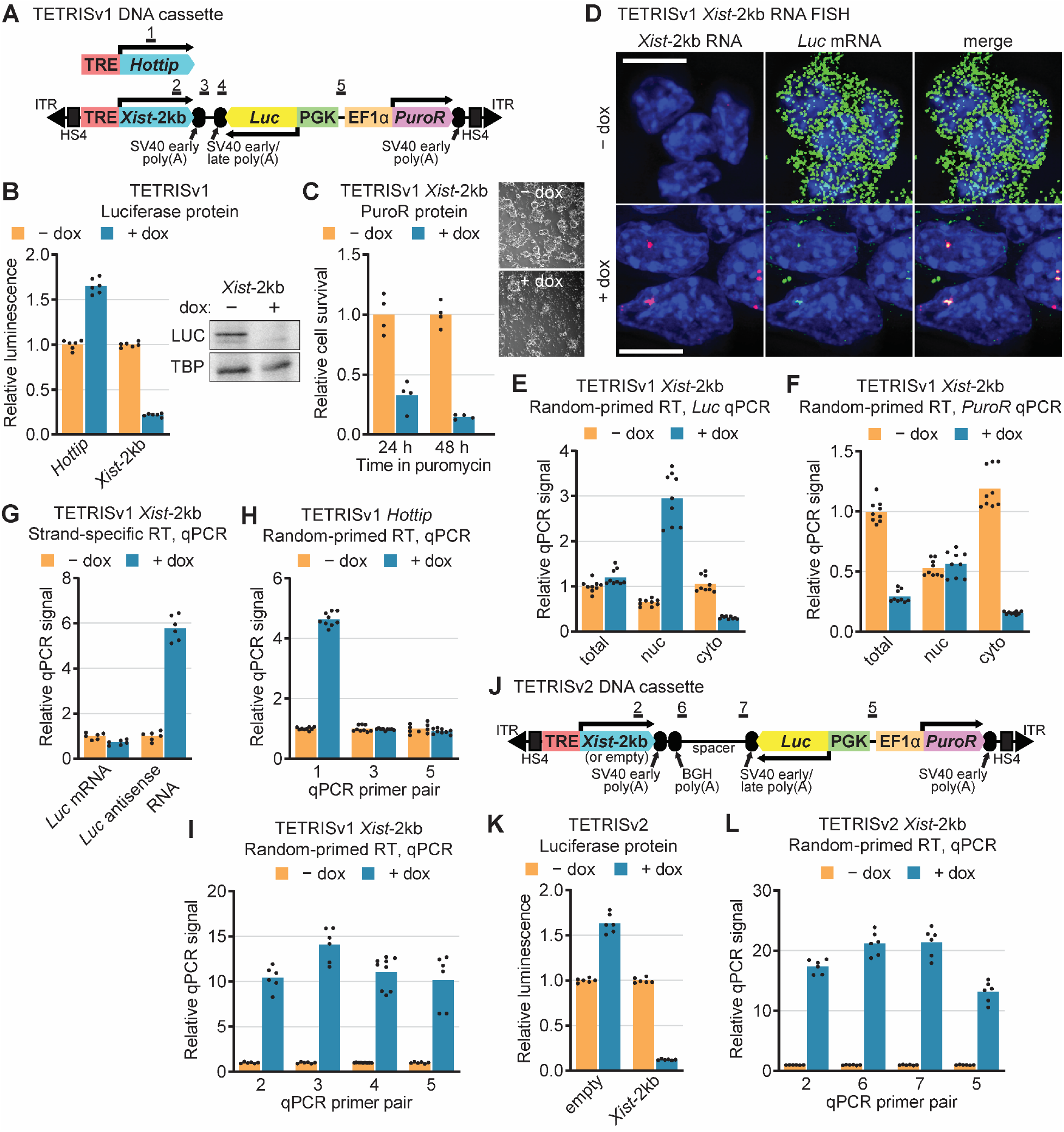
*Xist*-2kb represses expression of adjacent genes by reading through multiple polyadenylation sequences. (**A**) Schematics of TETRISv1 DNA cassettes. ITR, inverted tandem repeat recognized by piggyBac transposase; HS4, chicken β-globin insulator sequence; TRE, tetracycline response element (doxycycline-inducible promoter); *Xist*-2kb, nucleotides 1-2016 of mouse *Xist*; SV40, simian virus 40 (polyadenylation sequences); *Luc*, firefly luciferase gene; PGK and EF1α, constitutive promoters; *PuroR*, puromycin resistance gene. Numbered bars indicate regions targeted by qPCR primer pairs. (**B**) Luminescence assay (left) and western blot (right) showing effects of *Hottip* and *Xist*-2kb induction on luciferase protein expression. (**C**) Relative survival of TETRISv1 *Xist*-2kb cells grown for 24 or 48 h in the presence of puromycin after 48 h of treatment with or without doxycycline. Representative images show differences in cell survival after 48 h in puromycin. (**D**) Stellaris single-molecule RNA FISH shows *Xist* RNA (red) and *Luc* mRNA (green) in TETRISv1 *Xist*-2kb cells. DAPI-stained nuclei are blue. Scale bar = 10 μm. (**E-F**) RNA was prepared from TETRISv1 *Xist*-2kb cells directly (total) or following subcellular fractionation. Reverse transcription was performed with a pool of random oligos, and quantities of RNA corresponding to the *Luc* (E) or *PuroR* (F) gene in each sample were measured via qPCR. See Supplementary Figure S1A for fractionation/loading controls. (**G**) Total RNA from TETRISv1 *Xist*-2kb cells was reverse-transcribed using strand-specific primers targeting either the *Luc* mRNA or RNA produced from transcription occurring in the opposite direction (*Luc* antisense RNA). Quantities of RNA in each sample were measured with the same qPCR primer pair (relative to the – dox average for each strand-specific RT). (**H**) Nuclear RNA from TETRISv1 *Hottip* cells was reverse transcribed with random primers, and doxycycline-dependent changes in RNA produced from the *Hottip* gene and from the *Hottip-Luc* and *Luc*-*PuroR* intergenic regions were measured via qPCR with the primer pairs shown in (A). (**I**) Analysis as in (H) but with nuclear RNA from TETRISv1 *Xist*-2kb cells. (**J**) Diagram of the TETRISv2 DNA cassette and regions targeted by qPCR primer pairs in (L). (**K**) Luminescence assay as in (B) but with TETRISv2 empty (no lncRNA cargo) or *Xist*-2kb cells. (**L**) Analysis as in (H-I) but with TETRISv2 *Xist*-2kb cells. For all panels, cells were treated with or without 1 μg/mL doxycycline for 48 h prior to assaying. Unless stated otherwise, numerical values are shown relative to the – dox average for each cell identity or qPCR primer pair, which is set to one. Dots represent individual technical replicate measurements from a minimum of biological duplicate experiments, and bars represent the average value. See Supplementary Table 1 for information regarding experimental replicates and Supplementary Table 2 for oligo sequences.

We next performed controls to verify whether expression of *Xist*-2kb caused a level of transcriptional silencing commensurate with the observed reduction in luciferase protein levels. Indeed, we found via single-molecule sensitivity RNA FISH that, upon *Xist*-2kb expression, the number of luciferase mRNA molecules was greatly reduced throughout the cell (Figure 1D). We next analyzed RNA in subcellular fractions via reverse transcription (RT) with random primers followed by quantitative PCR (qPCR). In response to *Xist*-2kb expression, cytoplasmic qPCR signal for the *Luc* gene was reduced, in agreement with RNA FISH (Figure 1D and E, Supplementary Figure S1A). However, in response to *Xist*-2kb expression, nuclear qPCR signal for the *Luc* gene increased, as if *Luc* had not been transcriptionally silenced (Figure 1E, Supplementary Figure S1A). Likewise, in response to *Xist*-2kb expression, cytoplasmic qPCR signal from the *PuroR* gene decreased while nuclear *PuroR* signal remained relatively constant, as if that gene too had not been silenced (Figure 1F).

To reconcile these seemingly conflicting results, we examined the possibility that transcription from the *Xist*-2kb transgene failed to terminate at its specified polyadenylation sequence, causing readthrough into the downstream *Luc* and *PuroR* genes. Transcriptional readthrough would be indistinguishable from genic transcription as measured by the random-primed RT-qPCR assays of Figure 1E and F and could explain the relative increase in nuclear RT-qPCR signal over the *Luc* and *PuroR* genes induced by expression of *Xist*-2kb. To test this hypothesis, we performed strand-specific RT using primers targeting either the *Luc* mRNA or the putative readthrough transcription product. Subsequent qPCR confirmed that *Xist*-2kb expression led to readthrough transcription over the body of the *Luc* gene (Figure 1G).

To further examine transcriptional readthrough in the TETRIS assay, we used random-primed RT and a series of qPCR primer pairs targeting multiple sites within the TETRIS cassette, including primer pairs located downstream of *Xist*-2kb’s SV40 early polyadenylation sequence (numbering per Figure 1A). Expression of the control lncRNA *Hottip* caused no change in qPCR signal downstream of the SV40 early polyadenylation sequence, confirming that the sequence is functional (Figure 1H). On the other hand, expression of *Xist*-2kb caused a 10-to-15-fold increase in qPCR signal between the lncRNA gene and the *Luc* gene (primer pairs 3 and 4) that was commensurate to the level of *Xist*-2kb induction (primer pair 2), consistent with robust transcriptional readthrough of the SV40 early polyadenylation sequence (Figure 1I). Additional qPCR targeting the region between the *Luc* and *PuroR* genes (primer pair 5) suggests that readthrough transcription originating from *Xist*-2kb extends through *Luc* and into the downstream *PuroR* gene (Figure 1I). Remarkably, in order for this latter scenario to occur, transcription originating from the *Xist*-2kb transgene would have had to continue through a second SV40 early polyadenylation sequence, which is encoded by the reverse complement of the SV40 late polyadenylation sequence that terminates transcription of the *Luc* gene (see (46); Figure 1A). Sanger sequencing confirmed that the polyadenylation sequences in the TETRIS *Xist*-2kb and *Hottip* cargo plasmids were identical (Supplementary Figure S1B). Thus, in the context of TETRIS, transcription of *Xist*-2kb appears to extend well beyond its expected polyadenylation site.

We examined whether transcriptional readthrough beyond *Xist*-2kb would occur in a second version of TETRIS (TETRISv2) in which two consecutive polyadenylation sequences (SV40 early and bovine growth hormone [BGH]) were placed downstream of *Xist*-2kb, followed by a 1.1-kb spacer that separates these polyadenylation sequences from the SV40 early/late polyadenylation sequence of the convergently oriented *Luc* gene (Figure 1J). As with the original version of TETRIS, we observed that in TETRISv2, induction of *Xist*-2kb reduced luciferase activity by ~90% (Figure 1K). Even in this second vector, upon *Xist*-2kb induction, we observed robust RT-qPCR signal in the region between *Xist*-2kb and *Luc* (primer pairs 6 and 7) and in the region between *Luc* and *PuroR* (primer pair 5), which is located approximately 6.3 kb downstream of the transcription start site of the *Xist*-2kb gene (Figure 1L). Therefore, for *Xist*-2kb but not controls, transcription appeared to read through three near-consecutive polyadenylation sequences: SV40 early, BGH, and a second SV40 early sequence at the 3′ end of the *Luc* gene.

### Transcriptional readthrough by *Xist*-2kb requires Repeat A and downstream sequence

Given our results, we reasoned that *Xist*-2kb likely contains sequences that promote transcriptional readthrough of downstream polyadenylation sequences. Our prior data showed that in the context of the original TETRIS assay (TETRISv1), repression of luciferase activity by *Xist*-2kb required the same sequence elements within Repeat A that are required by full-length *Xist* to induce chromosome-level transcriptional silencing: its GC-rich repeats but not its U-rich linkers (4,36). We therefore investigated whether transcriptional readthrough induced by *Xist*-2kb required these GC-rich repeats (Figure 2A). Expression of a mutant *Xist*-2kb lacking the GC-rich repeats of Repeat A displayed reduced levels of transcriptional readthrough relative to wild-type *Xist*-2kb (Figure 2B), as well as reduced steady-state levels upon induction by doxycycline (~2-fold induction in Figure 2B vs. ~10-fold induction in Figure 1I). Our previous work also showed that in TETRISv1, repression of luciferase activity by *Xist*-2kb was attenuated by deletion of a ~750 nt region immediately downstream of Repeat A that contains elements predicted to form stable structures (“ss234”) (Figure 2A; (14,36)). Accordingly, we found that deleting this region almost completely prevented transcriptional readthrough despite a ~10-fold induction of the mutant *Xist-*2kb (Figure 2C). Lastly, deletion of Repeat A and the ss234 element together (“ΔrA234”, Figure 2A) reduced the levels of *Xist*-2kb induction and readthrough transcripts (Figure 2D). We conclude that transcriptional readthrough induced by *Xist*-2kb requires Repeat A as well as a ~750 nt downstream region.

**Figure 2.**
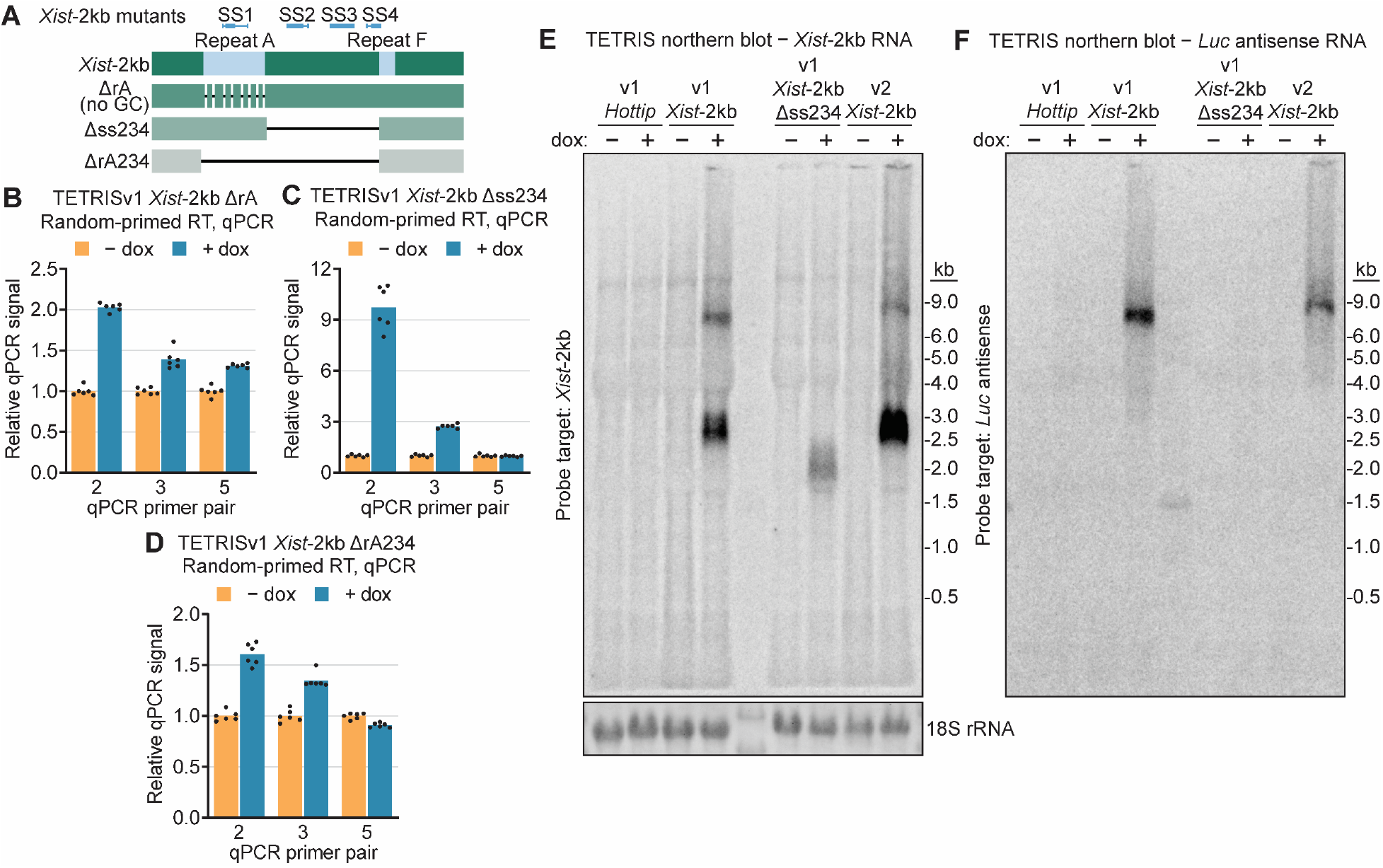
Transcriptional readthrough by *Xist*-2kb requires Repeat A and downstream sequence. (**A**) Diagram of *Xist*-2kb showing the location of Repeat A, Repeat F, stably structured elements (SS; (14)), and mutants that reduce repressive activity in TETRIS (36). Sequences of each mutant can be found in (36). The deleted ss234 region corresponds to nt 733-1474 of *Xist*, chromosomal coordinates chrX:103481759-103482500 in mm10. (**B-D**) Nuclear RNA from cells harboring TETRISv1 *Xist*-2kb deletion mutants was reverse-transcribed with random primers and analyzed by qPCR with primer pairs targeting regions shown in Figure 1A. Numerical values are shown relative to the – dox average for each qPCR primer pair, which is set to one. Dots represent individual technical replicate measurements, and bars represent the average value. (**E-F**) Northern blot analysis of total RNA prepared from cells harboring the indicated TETRIS expression cassettes. Following membrane transfer, ethidium-stained RNA was imaged to visualize evenness of loading (18S rRNA in (E)) and migration of RNA size markers. Membrane was probed first with a single-stranded DNA oligo targeting *Luc* antisense RNA (F), stripped and imaged to ensure removal of signal (not shown), and re-probed with a single-stranded DNA oligo targeting *Xist*-2kb (E). See Supplementary Figure S2 for additional images. For all panels, cells were treated with or without 1 μg/mL doxycycline for 48 h prior to assaying. See Supplementary Table 1 for information regarding experimental replicates and Supplementary Table 2 for oligo sequences.

To validate these results and observe the lengths of *Xist*-2kb readthrough transcripts, we analyzed RNA from untreated and doxycycline-treated TETRIS cells by northern blot, using single-stranded DNA probes targeting either *Xist*-2kb or RNA antisense to the luciferase mRNA product (Figure 2E and F). In both TETRISv1 and TETRISv2, 2349 bp separates *Xist*-2kb’s transcriptional start site from its expected cleavage site in the downstream SV40 early polyadenylation sequence (47). Accordingly, the *Xist*-2kb probe detected a prominent band around 2.5 kb in doxycycline-induced samples, consistent with cleavage at this site and the addition of a poly(A) tail (Figure 2E, Supplementary Figure S2B). However, distinct bands between the 6-kb and 9-kb size markers were also detected. These bands suggest a portion of readthrough transcripts terminate at the SV40 early polyadenylation sequence at the end of the *PuroR* gene, 7166 bp away from *Xist*-2kb’s transcriptional start site in TETRISv1 and 7735 bp away in TETRISv2. The higher-molecular-weight products were also detected with the *Luc* antisense-targeting probe, confirming that these RNAs contain both *Xist*-2kb sequence and sequence antisense to *Luc* mRNA (Figure 2F). As controls, *Hottip* and *Xist*-2kb lacking the ss234 region were detected with sequence-specific probes only at their expected sizes (Figure 2E and Supplementary Figure S2A). Moreover, the *Luc* antisense-targeting probe did not detect readthrough transcription products with *Hottip* or *Xist*-2kb Δss234 (Figure 2F), confirming efficient cleavage and polyadenylation of these lncRNAs. Thus, sequences within *Xist*-2kb harbor antiterminator activity that can suppress downstream 3′-end processing at two strong and commonly used polyadenylation sequences, SV40 early and BGH, resulting in robust transcriptional readthrough into downstream sequences.

### *Xist*-2kb induces mild levels of local transcriptional silencing in a SPEN-independent manner

The previous experiments unexpectedly revealed transcriptional readthrough as a confounding factor in our ability to determine whether *Xist*-2kb causes transcriptional silencing of adjacent reporter genes in the context of TETRIS. It was therefore unknown to what extent the ~80% repression of luciferase protein activity in TETRIS depended on transcriptional readthrough by the *Xist*-2kb transgene. Strand-specific RNA FISH in Figure 1D showed a loss in *Luc* mRNA signal throughout the cell but also suggested that some portion of *Luc* transcripts remain at their site of transcription in the nucleus. To investigate this result further, we performed strand-specific RT followed by qPCR on RNA extracted from cytoplasmic and nuclear fractions (Supplementary Figure S3A). This assay demonstrated that while expression of *Xist*-2kb caused a decrease in cytoplasmic levels of strand-specific *Luc* mRNA signal, nuclear levels were essentially unaffected (Supplementary Figure S3B). Thus, the nucleocytoplasmic ratio of *Luc* mRNA was shifted toward the nucleus (Supplementary Figure S3C). These data suggest that, although expression of the *Xist*-2kb transgene in a convergent orientation may impair *Luc* mRNA release from chromatin and nuclear export, *Xist*-2kb does not transcriptionally silence the *Luc* gene completely, as might have been expected (4,24,27). As a result, *Xist*-2kb’s ability to repress luciferase protein levels in TETRIS appears to be a direct consequence of transcriptional readthrough into the *Luc* gene.

To further address whether *Xist*-2kb functions as a transcriptional repressor, we designed a new version of TETRIS (TETRISv3) in which transcriptional readthrough would not affect reporter gene expression. To do this, we excised the doxycycline-inducible lncRNA expression cassette from TETRISv2, re-circularized the vector, and re-cloned the lncRNA expression cassette downstream of and in the same orientation as the *PuroR* gene, such that induction of *Xist*-2kb would not cause any readthrough transcripts to traverse either the *Luc* or *PuroR* genes (Figure 3A). In TETRISv3, expression of *Xist*-2kb caused only a ~2-fold reduction of luciferase protein activity relative to the empty vector control (Figure 3C and D). RT-qPCR indicated that the ~2-fold repression of luciferase protein activity in TETRISv3 was coincident with a ~2-fold reduction of nuclear *Luc* mRNA levels, indicating that *Xist*-2kb induced a mild amount of transcriptional silencing in this setting (Figure 3E). Similarly weak levels of transcriptional silencing were observed with *PuroR* mRNA (Figure 3F), and the nucleocytoplasmic ratios of both *Luc* and *PuroR* mRNAs were unchanged by *Xist*-2kb expression (Figure 3G and H).

**Figure 3.**
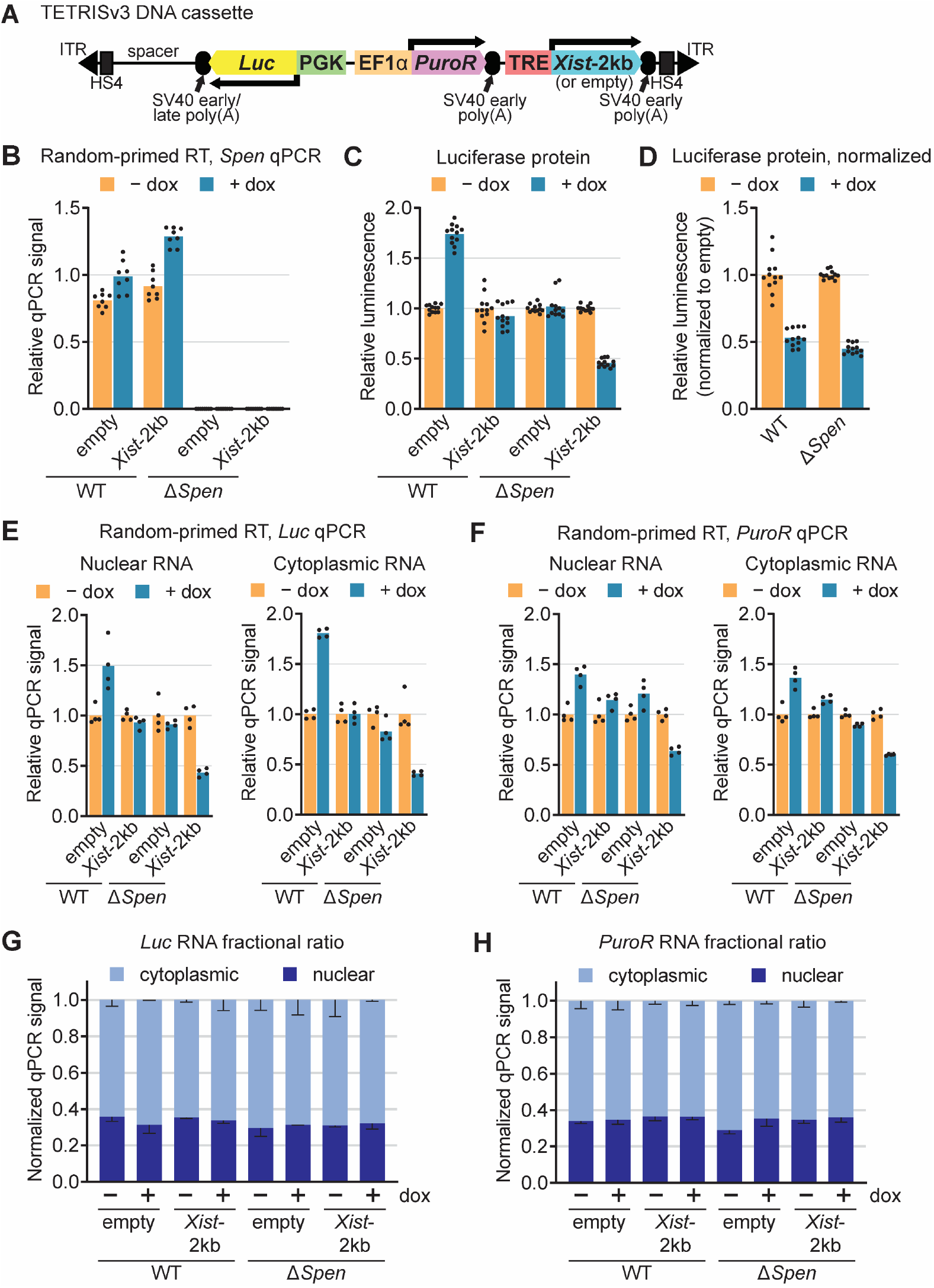
*Xist*-2kb induces mild levels of local transcriptional silencing in a SPEN-independent manner. (**A**) Diagram of TETRISv3 DNA cassette. (**B)** RT-qPCR showing levels of *Spen* mRNA in wild-type and Δ*Spen* cells harboring the TETRISv3 empty and *Xist*-2kb cassettes. See Supplementary Figure S4 for additional characterization of Δ*Spen* cells. (**C**) Luminescence assay showing relative luciferase protein levels in TETRISv3 empty and *Xist*-2kb cells treated with or without doxycycline for two days. (**D**) The *Xist*-2kb values in (C) are shown normalized to the corresponding empty values. (**E-F**) Random-primed reverse transcription of nuclear and cytoplasmic RNA from each cell line was followed by qPCR targeting either luciferase (E) or *PuroR* (F). (**G-H**) The sum of nuclear and cytoplasmic qPCR signal for each cell line in (E-F) was normalized to one to depict nucleocytoplasmic ratios. Dots represent individual technical replicate measurements, and bars represent the average value. See Supplementary Table S1 for information regarding experimental replicates and Supplementary Table S2 for oligo sequences.

Because the protein SPEN is required for transcriptional silencing induced by full-length *Xist* (10,23–28), we sought to determine whether SPEN was required for the mild levels of transcriptional silencing induced by the *Xist*-2kb transgene in TETRISv3. We used CRISPR to delete ~40kb of the *Spen* gene in ESCs, including its major RNA-binding domains (as done in (28); Supplementary Figure S4A). This deletion is known to cause complete failure of XCI and is expected to comprise a null mutant (28). The deletion was confirmed in select clones by PCR of genomic DNA (Supplementary Figure S4B) and RT-qPCR, which showed the expected loss of *Spen* mRNA expression in the deleted region (Figure 3B, Supplementary Figure S4C). Using these *Spen* knockout ESCs, we then generated TETRISv3 cells inducibly expressing *Xist*-2kb or an empty expression cassette. In two independent SPEN knockout clones, we found that the levels of *Luc* silencing induced by *Xist*-2kb relative to empty vector control in TETRISv3 were commensurate to those induced in wild-type ESCs (Figure 3C and D). Thus, in the context of TETRIS, the ~2-fold transcriptional silencing induced by the *Xist*-2kb transgene is SPEN-independent.

### *Xist*-2kb binds SPEN and RBM15, but these proteins are not required for transcriptional readthrough

We next sought to determine whether *Xist*-2kb expressed in TETRIS interacts with proteins known to bind this region in the context of full-length *Xist*. To answer this question, we first used an antibody-independent method called capture hybridization analysis of RNA targets coupled with mass spectrometry (CHART-MS), in which biotinylated DNA oligonucleotides are used to capture an RNA of interest and its bound proteins from nuclear extracts prepared from formaldehyde-crosslinked cells (48,49). Specifically, we performed CHART-MS to compare the proteins associated with wild-type *Xist*-2kb to those associated with a nonfunctional mutant version of *Xist*-2kb lacking Repeat A and the ~750-nt ss234 region (ΔrA234, Figure 2A). We identified 22 proteins that associated with wild-type *Xist*-2kb in two independent replicate experiments but not with the ΔrA234 mutant *Xist* in the single replicate performed (Supplementary Table S3). The full list of proteins retrieved in each of the three experiments is shown in Supplementary Table S4, and Scaffold output data are available as Supplementary File 1. Two of the proteins enriched specifically by *Xist*-2kb were SPEN and RBM15, both of which associate with Repeat A in the context of full-length *Xist* (10,23–28,50). Also identified were proteins known to play roles in splicing and RNA export, including U2AF2, DHX15, SRSF6, NXF1, and SNRNP200, consistent with previous studies linking Repeat A to RNA processing (Supplementary Table S3; (7,16,20–22)). Using a formaldehyde-based RNA immunoprecipitation protocol (45), we confirmed that SPEN and RBM15 associated with RNA produced from the *Xist*-2kb transgene in a Repeat A-dependent manner (Figure 4A and B).

**Figure 4.**
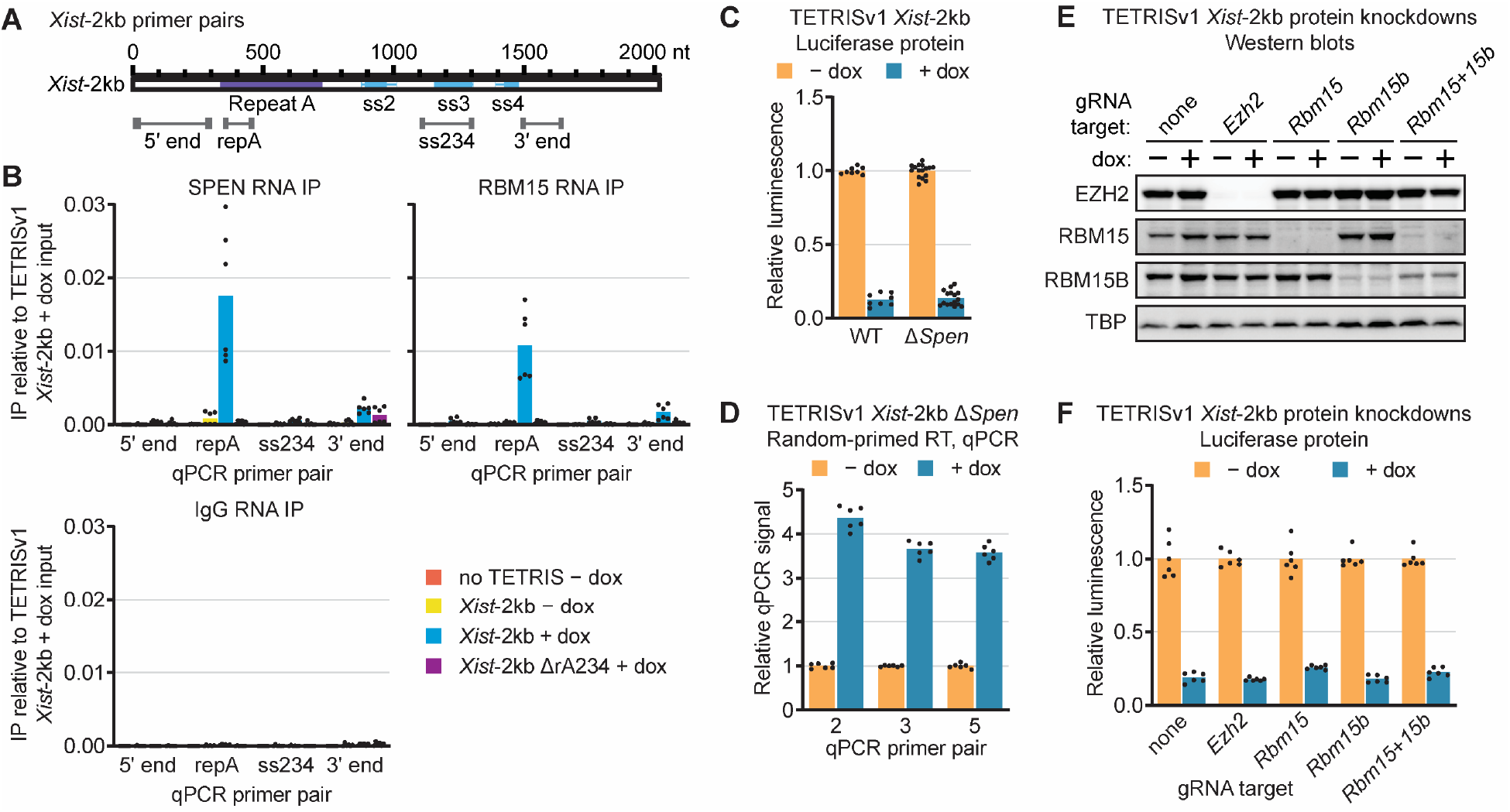
*Xist*-2kb binds SPEN and RBM15, but these proteins are not required for transcriptional readthrough. (**A**) Diagram of *Xist*-2kb showing locations of qPCR primer pairs. (**B**) Four different cell identities (cells without TETRIS cargo, TETRISv1 *Xist*-2kb cells treated with or without doxycycline, and TETRISv1 *Xist*-2kb ΔrA234 cells treated with doxycycline) were used for RNA immunoprecipitation with SPEN, RBM15, or IgG control antibodies, followed by random-primed RT and qPCR with primer pairs shown in (A). Values are relative to *Xist*-2kb + dox input. (**C**) Luminescence assay in wild-type and Δ*Spen* cells harboring TETRISv1 *Xist*-2kb cargo. (**D**) Nuclear RNA from Δ*Spen* cells harboring TETRISv1 *Xist*-2kb cargo was reverse-transcribed with random primers and analyzed by qPCR with primer pairs targeting regions shown in Figure 1A. (**E**) Western blots demonstrating successful CRISPR-Cas9 targeting of *Ezh2*, *Rbm15*, *Rbm15b*, and both *Rbm15* and *Rbm15b* in cells harboring TETRISv1 *Xist*-2kb cargo. (**F**) Luminescence assays with CRISPR knockdown TETRISv1 *Xist*-2kb cells shown in (E). See Supplementary Figure S4 for characterization of Δ*Spen* cells and for locations of CRISPR gRNAs. Dots represent individual technical replicate measurements, and bars represent the average value. See Supplementary Table S1 for information regarding experimental replicates and Supplementary Table S2 for oligo sequences.

Because transcriptional readthrough induced by *Xist*-2kb required Repeat A (Figure 2B,D), we hypothesized that transcriptional readthrough might also require the Repeat A-binding proteins SPEN or RBM15. TETRISv1 assays performed in two independent SPEN deletion lines (Supplementary Figure S4) showed that SPEN deletion had no effect on luciferase repression induced by *Xist*-2kb (Figure 4C). Given that luciferase repression induced by *Xist*-2kb in TETRISv1 is largely the result of transcriptional readthrough, this result implies that SPEN is not required for transcriptional readthrough. Accordingly, subsequent RT-qPCR assays provided direct evidence for this notion (Figure 4D), although we did observe that in the absence of SPEN, lower levels of *Xist*-2kb were induced than in SPEN-expressing cells (~4-fold in Figure 4D vs. ~10-fold in Figure 1I). To examine the requirement of RBM15 for transcriptional readthrough induced by *Xist*-2kb, polyclonal cell populations were generated that carried sgRNAs targeting a doxycycline-inducible Cas9 to numerous locations in exons of the *Rbm15* gene, leading to depletion of the protein product (Figure 4E, Supplementary Figure S4E). RBM15B, a paralog of RBM15, was also depleted individually and in combination with RBM15 (Figure 4E, Supplementary Figure S4F). EZH2, which has also been shown to associate with Repeat A (6,9,11), was also depleted (Figure 4E, Supplementary Figure S4D). Relative to control cells that expressed a non-targeting sgRNA, TETRIS luminescence assays in RBM15, RBM15B, RBM15+RBM15B, and EZH2 knockdown cell lines showed no hindrance of luciferase protein repression by *Xist*-2kb (Figure 4F). Therefore, SPEN, RBM15, RBM15B, and EZH2 do not appear to be required for transcriptional readthrough induced by *Xist*-2kb.

### As single-copy transgenes, 5′ fragments of *Xist* cause transcriptional readthrough beyond their polyadenylation sites

Under the conditions we used to make standard TETRIS cell lines, approximately five copies of the TETRIS DNA cassette are randomly inserted into the genome of each cell that survives the selection process (36). Thus, while TETRIS is suitable to determine the local extent of silencing induced by lncRNAs, it cannot report on long-distance silencing. We therefore sought to determine the extent to which *Xist*-2kb and other *Xist* transgenes could promote transcriptional readthrough and long-distance transcriptional silencing by inserting the transgenes as a single copy into a defined chromosomal locus.

To this end, we established a recombinase-mediated cassette exchange (RMCE) system in the *Rosa26* locus of chromosome 6 (51). We created a *Rosa26*-targeting vector that contained a lox66 site, a *PuroR*-*ΔTK* fusion gene (52), and a lox2272 site followed by a BGH polyadenylation sequence (Supplementary Figure S5A). This targeting construct was electroporated into F1-hybrid, male ESCs that were derived from a cross between C57BL/6J (B6) and CAST/EiJ (Cast) mice that enable allelic analysis of gene expression. Southern blot was used to confirm insertion of the construct into the correct locus on the B6 allele of selected clones (Supplementary Figure S5B). In parallel, we created a cargo vector that contained a lox71 site, a lncRNA-expression cassette driven by a doxycycline-inducible promoter, a tandemly oriented and constitutively expressed hygromycin B resistance gene (*HygroR*) lacking a polyadenylation sequence, and a lox2272 site (Supplementary Figure S5C). Electroporation of the cargo vector along with Cre recombinase into our F1-hybrid RMCE cells, followed by positive selection with hygromycin B and negative selection with ganciclovir, generated a small number of surviving clones that harbored cargo vectors inserted in the desired orientation in *Rosa26* (Supplementary Figure S5D; not shown).

We employed this RMCE system to create four separate ESC lines that expressed different versions of inducible *Xist* transgenes from *Rosa26* (Figure 5A): one line expressed the *Xist*-2kb transgene, another expressed full-length *Xist* from its endogenous DNA sequence (including its introns and natural polyadenylation sequence), another expressed the first 5.5 kb of *Xist* (“*Xist*-5.5kb”), which includes the Repeat B and Repeat C domains of *Xist* known to recruit Polycomb repressive complex 1 (PRC1 (15,16,19,20)), and a final line expressed *Xist*-2kb fused to the final two exons of *Xist* (“*Xist*-2kb+6,7”, which includes the intron between exons 6 and 7). The final exon of *Xist* includes Repeat E and is essential for proper *Xist* localization (17,18,53). As a control, we created an ESC line that underwent recombination but lacked any *Xist* insertion (“empty”). We then used piggyBac-mediated transgenesis to insert the *rtTA* gene into select clones of each genotype in order to allow doxycycline-inducible expression of each cargo RNA. RNA FISH confirmed that RNA produced from each *Xist* transgene was doxycycline-inducible and remained localized in the nucleus, presumably surrounding its site of transcription (Figure 5B). We note that despite several attempts, we were unsuccessful in cloning an *Xist*-5.5kb construct that contained all ~36 repeats in Repeat B; the construct used for this study contained ~14 repeats (Figure 5A and Supplementary Figure S6A).

**Figure 5.**
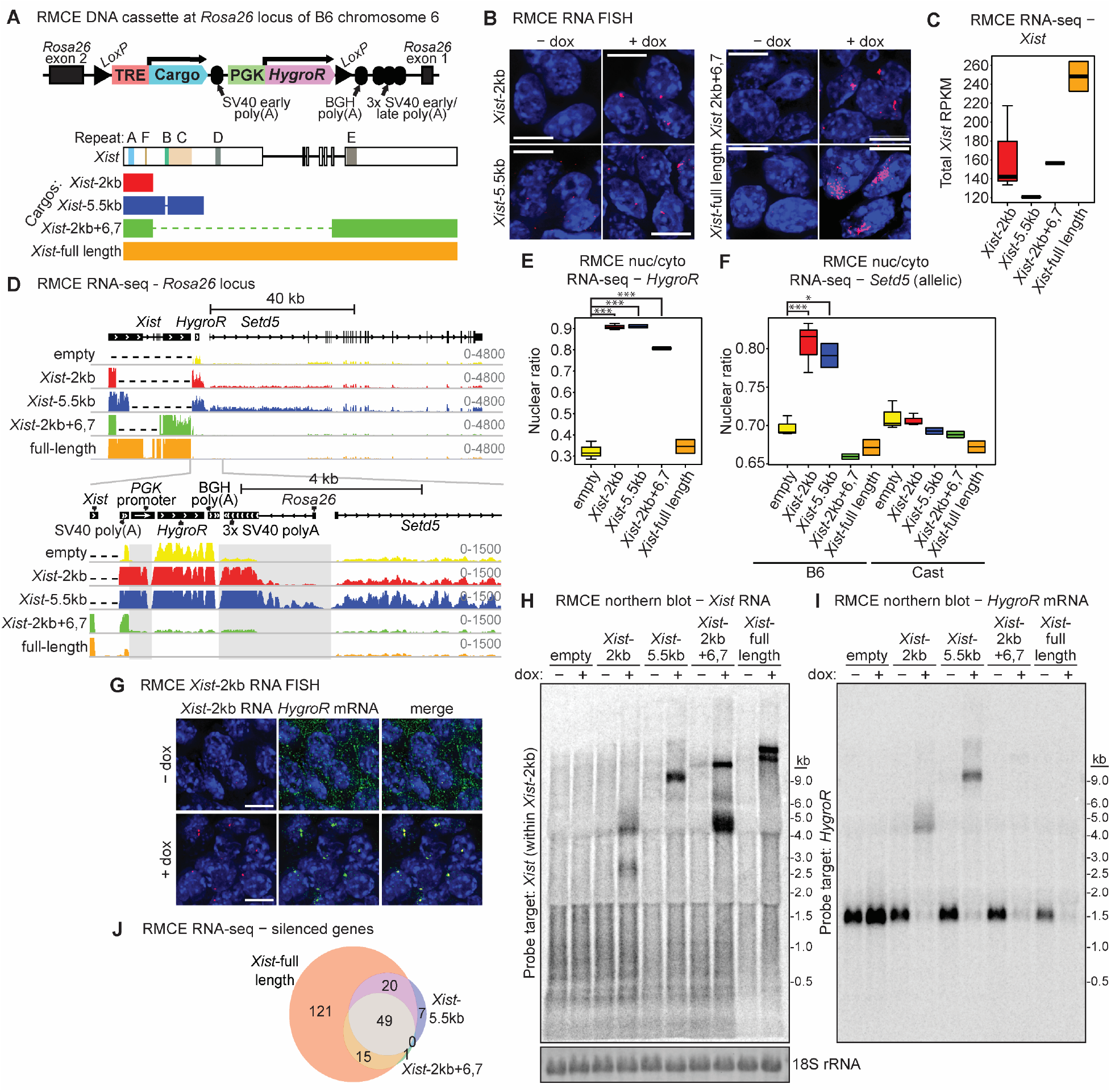
As single-copy transgenes, 5′ fragments of *Xist* cause transcriptional readthrough beyond their polyadenylation sites. (**A**) Diagram of recombination-mediated cassette exchange (RMCE) expression cassette inserted into the *Rosa26* locus on the B6 allele of chromosome 6 in B6/Cast F1 hybrid ES cells. Full-length *Xist* and three mutant transgene cargos were inserted into this expression cassette under a doxycycline-inducible promoter. *Xist*-5.5kb is missing a portion of Repeat B (Supplementary Figure S6A). *HygroR*, constitutively expressed hygromycin resistance gene. (**B**) Stellaris single-molecule RNA FISH for *Xist* RNA (red) in cells expressing *Xist*-2kb, *Xist*-5.5kb, *Xist*-2kb+6,7 or full-length *Xist* inserted at *Rosa26*, treated with or without doxycycline. DAPI-stained nuclei are blue. Scale bar = 10 μm. (**C**) *Xist* expression in each cell line following 3 d doxycycline treatment, determined via RNA-seq. Reads per kilobase per million (RPKM) values were calculated using reads aligned to the endogenous *Xist* locus divided by length of inserted transcript in kb divided by total aligned reads in millions for each dataset. (**D**) RNA-seq reads were aligned to a custom-made IGV build containing the single-insertion *Xist*/*HygroR* cassette within the *Rosa26* locus on chromosome 6. Top panel depicts this cassette upstream of the endogenous *Setd5* gene. Dashed lines within the *Xist* locus represent sequences not present in each version of the RMCE cassette. The bottom panel shows a closer view of RNA-seq reads extending beyond the SV40 polyadenylation sequence at the 3′ end of *Xist*, with regions of clear transcriptional readthrough highlighted in grey. (**E**) Nuclear ratio of reads mapping to *HygroR* following doxycycline treatment. (**F**) Nuclear ratio of allele-specific reads mapping to *Setd5* following doxycycline treatment. Note the difference in y-axis compared to (E). ***P < 0.0001; *P < 0.01, Tukey HSD post-hoc analysis of significant differences by ANOVA. (**G**) Stellaris single-molecule RNA FISH showing *Xist*-2kb RNA (red) and *HygroR* mRNA (green) in cells with *Xist*-2kb inserted at *Rosa26*. DAPI-stained nuclei are blue. Scale bar = 10 μm. (**H-I**) Northern blot analysis of RNA prepared from cells harboring the indicated RMCE expression cassettes. Following membrane transfer, ethidium-stained RNA was imaged to visualize evenness of loading (18S rRNA in (H)) and migration of RNA size markers (Supplementary Figure S7). Membrane was probed first with a single-stranded DNA oligo targeting *HygroR* (I), stripped and imaged to ensure removal of signal (not shown), then re-probed with a single-stranded DNA oligo targeting *Xist* within *Xist*-2kb (H). (**J**) Venn diagram showing overlaps of significantly repressed genes in *Xist*-5.5kb, *Xist*-2kb+6,7 and full-length *Xist* cells. See Supplementary Table S5 for RNA-seq gene expression data and Supplementary Figures S5-S9 for additional information related to Figure 5. See Supplementary Table S1 for information regarding experimental replicates and Supplementary Table S2 for oligo sequences.

To determine whether the *Xist* transgenes caused transcriptional readthrough of their downstream SV40 early polyadenylation sequence as well as to determine their effects on the expression of genes across chromosome 6, we induced the expression of each *Xist* transgene for three days and performed RNA-seq using RNA purified from cytoplasmic and nuclear fractions. In parallel, we treated empty-cargo ESCs with doxycycline for three days and sequenced RNA purified from cytoplasmic and nuclear fractions. Expression of the *Xist* transgenes was verified by examining reads mapping to the endogenous *Xist* locus (which is not expressed in these cells; Supplementary Figure S6B), and total counts were used to calculate RPKM values for each *Xist* transgene. Consistent with RNA FISH data in Figure 5B, RNA produced from each *Xist* transgene localized in the nucleus (Supplementary Figure S6C), and expression levels were only slightly lower for the *Xist* hypomorphs than for full-length *Xist* (Figure 5C).

We examined our RNA-seq data for evidence of *Xist*-dependent transcriptional readthrough in the RMCE setting. We aligned our RNA-seq data to an *in-silico* genome build that contained the transgenic vector features as they were inserted into *Rosa26* and then visualized the aligned reads in the Integrative Genomics Viewer (IGV; (54)). Consistent with our results above, when *Xist*-2kb, but not empty-cargo control, was induced, we observed robust transcriptional readthrough of the SV40 early polyadenylation sequence; this readthrough extended through the *PGK* promoter that drives expression of *HygroR*, through the BGH polyadenylation sequence at the 3′ end of *HygroR*, and through the bidirectionally functional triple-SV40-polyadenylation sequences designed to terminate expression of *Rosa26* (Figure 5D, regions shaded in grey). Small amounts of readthrough transcription appeared to extend through the *Rosa26* promoter and into the *Setd5* gene located just downstream. Readthrough transcription appeared more robust in cells expressing *Xist*-5.5kb but was barely apparent in cells expressing *Xist*-2kb+6,7 and was not apparent in cells expressing full-length *Xist*. In agreement with these data and the nuclear localization of the *Xist* transgenes (Supplementary Figure S6C), induction of *Xist*-2kb and *Xist*-5kb caused a dramatic shift of *HygroR* reads to the nucleus, suggesting the production of chimeric, chromatin-bound readthrough RNAs containing both *Xist* transgene and *HygroR* sequence (Figure 5E). Expression of *Xist*-2kb and *Xist*-5kb also caused mild, yet significant shifts of syntenic *Setd5* reads to the nucleus (Figure 5F), consistent with low-level readthrough extending into this gene. Inducing *Xist*-2kb+6,7 caused a less-pronounced nuclear shift in *HygroR* reads and no significant shift in *Setd5* reads, consistent with *Xist*-2kb+6,7 causing only trace amounts of transcriptional readthrough. Induction of full-length *Xist* had no effect on the nucleocytoplasmic distribution of *HygroR* or *Setd5* reads, reflecting efficient transcriptional termination of this transgene. Remarkably, RNA FISH using probes designed to detect RNA produced from the *HygroR* gene gave an *Xist*-like staining pattern after induction of *Xist*-2kb, supporting the occurrence of readthrough transcription and indicating that chimeric *Xist*-*HygroR* transcripts associate with chromatin in *in cis*, in an *Xist*-like manner (Figure 5G).

To corroborate these results, we used northern blot analysis (Figure 5H and I, Supplementary Figure S7). Cleavage occurring at the SV40 early polyadenylation site downstream of *Xist*-2kb would be expected to produce a 2261-nt RNA (excluding its poly(A) tail). About half of *Xist*-2kb signal migrated around 2.5 kb, suggesting some level of cleavage occurs at this site (Figure 5H). However, the other half of *Xist*-2kb signal migrated around 4 to 5 kb. This long RNA was also detected with a probe targeting *HygroR* (Figure 5I), suggesting that, in the RMCE setting, about half of *Xist*-2kb transcripts do not terminate at the expected polyadenylation site but instead continue downstream through the *HygroR* gene. Termination of the *Xist*-2kb-*HygroR* chimeric transcript at the BGH polyadenylation site downstream of *HygroR* or at the three SV40 late polyadenylation sites further downstream would produce RNAs 4155, 4501, 4755, and 5009 nt in length (disregarding poly(A) tail length), in agreement with the molecular weight of the larger product observed via northern blot.

For *Xist*-5.5kb, cleavage at its SV40 early polyadenylation site would be expected 5598 nt downstream of its transcription start site. However, no product of this size was detected (Figure 5H). Instead, nearly all signal migrated at an approximate size of 9 kb. This signal was also detected with the *HygroR*-specific probe, indicating that nearly all *Xist*-5.5kb transcripts continued downstream into the *HygroR* gene (Figure 5I). The ~9-kb size estimated for this RNA slightly exceeds the lengths expected for cleavage at the BGH and SV40 late poly(A) sites (7492, 7838, 8092, and 8346 nt). For both *Xist*-2kb and *Xist*-5.5kb, smears of signal at even higher molecular weights are consistent with RNA-seq data suggesting readthrough into *Setd5* (Figure 5D). Additionally, upon expression of both *Xist*-2kb and *Xist*-5.5kb, the 1.5-kb *HygroR* mRNA signal was lost, suggesting that transcriptional readthrough from *Xist*-2kb and *Xist*-5.5kb interfered with transcription initiation at the *HygroR* promoter. These data indicate that *Xist*-2kb and *Xist*-5.5kb can robustly suppress downstream cleavage and polyadenylation to generate abundant transcriptional readthrough products that contain *HygroR* sequence. Moreover, our RNA-seq and RNA FISH data suggest that, rather than being exported to the cytoplasm, chimeric *Xist*-*HygroR* RNAs are retained at their sites of transcription in an *Xist*-like manner (Figure 5E and G).

In contrast to *Xist*-2kb and *Xist*-5.5kb, induction of *Xist*-2kb+6,7 produced an RNA band of length greater than 9 kb (Figure 5H), in agreement with its expected size of 10,079 nt (excluding poly(A) tail length). Inducing full-length *Xist* (expected size of 18,001 nt from CMV TSS to annotated endogenous 3′ end) resulted in a doublet band at high molecular weight (Figure 5H), which may reflect the production of two *Xist* isoforms due to alternative splicing within exon 7 or alternative polyadenylation (55,56). The same alternative RNA processing event may be responsible for generating a second product between 4 and 5 kb upon induction of *Xist*-2kb+6,7 (Figure 5H). Induction of *Xist*-2kb+6,7 and full-length *Xist* both caused strong reductions in 1.5-kb *HygroR* mRNA signal (Figure 5I). Only a very faint signal migrating around the size of *Xist*- 2kb+6,7 was detected with the *HygroR* probe, consistent with trace amounts of transcriptional readthrough suggested by RNA-seq (Figure 5D and E). No *HygroR* signal was detected at higher molecular weights upon expression of full-length *Xist*, consistent with efficient transcriptional termination at *Xist*’s natural polyadenylation sequence. Together, these results confirm a lack of robust transcriptional readthrough beyond *Xist*-2kb+6,7 and full-length *Xist* and suggest that the doxycycline-dependent loss in *HygroR* mRNA signal is due to repressive activity of the *Xist*- 2kb+6,7 and full-length *Xist* RNAs.

### Relative to longer *Xist* transgenes, *Xist*-2kb is severely deficient in its ability to induce long-distance transcriptional silencing

Next, we examined the extent to which the different *Xist* transgenes silenced gene expression on their syntenic chromosome (Supplementary Figure S8). In cells expressing full-length *Xist*, 2404 genes were differentially expressed relative to empty-cargo ESCs. On the B6 allele of chromosome 6 – the chromosome that harbors the *Xist* transgene at the *Rosa26* locus – 214 genes were differentially expressed (Supplementary Figure S8A). Of these, 205 genes shifted in the downward direction and 9 shifted in the upward direction, consistent with repression by full-length *Xist* (Supplementary Figure S8A). On the Cast allele of chromosome 6, only 34 genes changed, with similar numbers going up (21) and down (13; Supplementary Figure S8B). Throughout the genome, near-equal numbers of up- and down-regulated genes were detected (1012 and 1163, respectively). Thus, expression of full-length *Xist* caused chromosome-level repression of genes *in cis* and genome-wide changes in gene expression, the latter presumably being due to secondary effects.

Expression of *Xist*-5.5kb and *Xist*-2kb+6,7 also caused gene silencing, but at a reduced level relative to full-length *Xist*. *Xist*-5.5kb expression led to differential expression of 98 genes on the B6 allele of chromosome 6, 76 of which were shifted down (Supplementary Figure S8C). *Xist*- 2kb+6,7 expression led to differential expression of 88 genes on the B6 allele of chromosome 6, 65 of which were shifted down (Supplementary Figure S8E). The majority of genes repressed by these two transgenes were also repressed by full-length *Xist* (69 of 76 for *Xist-*5.5kb and 64 of 65 for *Xist*-2kb+6,7; Figure 5J). Loss of nuclear as well as cytoplasmic RNA-seq signal for these genes indicated that, like full-length *Xist*, the hypomorphic *Xist* transgenes induced transcriptional silencing (Supplementary Figure S9). In contrast, when comparing cytoplasmic RNA levels between *Xist*-2kb and empty-cargo ESCs, we only detected two differentially expressed genes genome-wide (excluding *Xist*), both on the B6 allele of chromosome 6. These genes were *Creld1*, located 406 kb downstream of the Rosa26 locus, and *Trh*, located ~20 Mb upstream (Supplementary Figure S8G). These data indicate that, relative to the longer *Xist* transgenes, *Xist*-2kb is severely deficient in its ability to induce long-distance transcriptional silencing. Deficient silencing by *Xist*-2kb in the RMCE setting is consistent with our observations using TETRIS (Figure 3 and Supplementary Figure S3) and with observations made previously (4,8,16).

## Discussion

Transcriptional silencing during XCI is thought to initiate when Repeat A recruits SPEN; SPEN recruitment, in turn, is thought to recruit and/or locally activate histone deacetylases and corepressor proteins to silence transcription over the X chromosome (10,23–28). Our data demonstrate that in mouse ESCs, recapitulating a SPEN-Repeat A interaction on chromatin is insufficient to induce robust local or long-distance transcriptional silencing by *Xist*, highlighting a gap in our understanding of the mechanism through which Repeat A functions to silence gene expression. We found that even though *Xist*-2kb associates with SPEN and localizes to chromatin in a manner that is indistinguishable from silencing-competent versions of *Xist*, it fails to induce robust transcriptional silencing, even of nearby genes. However, fusion of *Xist*-2kb to the next 3.5 kb of *Xist* (*Xist*-5.5kb), which in our construct included Repeat C and a portion of the essential PRC1-recruitment domain Repeat B (Figure 5A, Supplementary Figure S6A; (15,16,19,20)), or the fusion of *Xist*-2kb to the final two exons of *Xist* (*Xist-*2kb+6,7), which lacks a PRC1 recruitment domain but contains Repeat E and additional downstream sequence elements (14,17,53,57), both conferred near-equal transcriptional silencing capability in an isogenic context (Figure 5J, Supplementary Figures S7 and S8). Thus, in mouse ESCs, Repeat A is necessary for long-distance transcriptional silencing induced by *Xist*, but it is not sufficient. Silencing requires synergy between Repeat A and additional downstream sequence elements within *Xist*. It will be important to define the molecular mechanisms that underlie this synergy in future works.

During the course of our study, we also found, quite unexpectedly, that sequences within the 5′ end of *Xist* harbor transcriptional antiterminator activity that can cause transcriptional readthrough of consecutive strong polyadenylation sequences – specifically, the SV40 early and bovine growth hormone poly(A) sequences. These observations were made using two reductionist systems similar in nature to those previously employed to identify seminal aspects of *Xist* biology (4,8,10,14,16,19,23–28,32). In the context of the TETRIS assay, we found that the antiterminator activity of *Xist*-2kb required the GC-rich portions of Repeat A and a 742-nt region located just downstream that contains elements predicted to form stable structures (13,14,58). In the context of our RMCE system, *Xist*-2kb again displayed transcriptional antiterminator activity, while *Xist*-5.5kb displayed even stronger antiterminator activity, suggesting sequences downstream of *Xist*’s first two kilobases may enhance suppression of cleavage and polyadenylation. Longer, spliced *Xist* transgenes, namely *Xist*-2kb+6,7 and full-length *Xist*, terminated at expected sites, suggesting that the 5′ antiterminator activity loses its potency at longer range and/or the act of splicing dampens it.

It has been proposed that during the early stages of transcription, a sequence-non-specific mechanism suppresses premature 3′ end formation at polyadenylation sequences located near the 5′ end of genes (59). While a sequence-non-specific mechanism may exist, our observations show a pronounced sequence specificity – in TETRIS, transcriptional readthrough by *Xist*-2kb was abrogated, not enhanced, when elements were deleted that brought the polyadenylation sequences closer to the 5′ end of the transgene. Additionally, in TETRIS, transcriptional termination was efficient for the *Hottip* gene, which in our assays included an intron and produced a spliced RNA similar in length to *Xist*-2kb (Supplementary Figure S2A). Similarly, in the RMCE setting, suppression of cleavage and polyadenylation was more pronounced with *Xist*-5.5kb than with the shorter *Xist*-2kb transgene; moreover, induced transcription of the “empty-cargo” construct did not cause transcriptional readthrough into the *HygroR* gene despite the TRE promoter being in closer proximity to the SV40 early polyadenylation sequence. Therefore, it is likely that specific elements within the 5′ end of the *Xist* RNA actively suppress downstream cleavage and polyadenylation. In turn, these elements may promote production of full-length *Xist* in its endogenous context. Indeed, PolyA_SVM predicts that *Xist* contains strong polyadenylation sites 2497 and 6347 nt from its 5′ end (Supplementary Figure S10; (60,61)). Suppressing 3′ end formation at these sites may be a crucial purpose of the antiterminator activity we describe. Sequence-specific, RNA-mediated transcriptional antitermination is well-documented in bacteria (62), so it is conceivable that similar antiterminator elements exist in mammalian RNAs, even beyond *Xist*.

Mechanistically, *Xist*’s 5′ antiterminator activity may be related to a process called telescripting, in which the U1 snRNP, independent of its role in splicing, prevents premature termination of pre-mRNAs at cryptic cleavage and polyadenylation sites, particularly in long genes (63–65). While the 5′ end of *Xist* does not harbor obvious U1 snRNA binding sites (66), we and others have found that Repeat A associates with protein components of the U1 and other snRNPs (Supplementary Table S3; (7,16,21)). Moreover, several proteins that were identified as *Xist* cofactors in recent genetic and proteomic screens – namely NXF1, SRRT, and SCAF4 – have known roles in suppressing premature cleavage and polyadenylation (10,21,26,67–69). Future work will determine whether these proteins or others mediate *Xist*’s antiterminator activity.

We speculate that an antiterminator function may be the major reason why deletion of Repeat A or its relocation further downstream can cause such a dramatic reduction in the abundance of full-length *Xist* (7,20,27,29,31–35). Premature polyadenylation can repress transcription from nearby promoters as well as cause transcriptional attrition over the length of a gene (65,70). Moreover, the *Xist* RNA associates with multiple proteins that function to repress transcription – these include SPEN, which is recruited by Repeat A; PRC1, which is recruited by Repeats B and C; and PRC2, which is presumably recruited, directly and indirectly, by multiple regions in *Xist* including Repeats A and E (6,9–12,15–20). Prior work has shown in mouse embryonic fibroblasts and cardiomyocytes that certain genes can attenuate their own transcription by recruiting PRC2 through intronic RBFOX2-binding elements embedded in their nascent pre-mRNAs (71). Thus, given the large number of repressive complexes known to associate with the *Xist* RNA, precedent supports the notion that the *Xist* gene would require a mechanism to evade self-silencing. Moreover, *in vitro*, SPEN has no obvious preference for binding wild-type Repeat A over non-functional versions of Repeat A whose internal secondary structures have been minimally disrupted by point mutations or a two-nucleotide deletion (28). It is plausible that a need to derive an antiterminator activity could have been a driving force in the evolution of Repeat A, enabling it to recruit proteins in addition to SPEN whose functions are to suppress premature cleavage and polyadenylation and allow robust expression of an RNA capable of chromosome-wide silencing.

## Data availability

All genomic data, including raw sequencing files and processed data files, are available at GEO accession number GSE120197. Genomic data are also available in wiggle tracks at UCSC genome browser: http://genome.ucsc.edu/s/davidlee/rosa26_xist_fractions_lee_2019.

## Supplementary data and methods

Supplementary data and methods are available at *NAR* online.

## Author contributions

J.B.T., D.M.L., and J.M.C. conceived the study; D.O.C., S.R.B., and J.M.C. designed the RMCE strategy; J.B.T., D.M.L., R.E.C., S.O.K., K.I., M.D.S., S.R.B., and J.M.C. designed and performed experiments, and J.B.T., D.M.L., and J.M.C. wrote the paper.

## Acknowledgements

We would like to thank the editors at *NAR* for making this study possible. We also thank D. Ciavatta, N. Hathaway, M. Magnuson, and T. Magnuson for reagents, Y. Xu for contributions to Figure 1B, the UNC Microscopy Services Laboratory, and the Mass Spectrometry Facility at the University of Massachusetts Medical School.

