## Supplementary Methods and Figures for "Elements at the 5′ end of *Xist* harbor SPEN-independent transcriptional antiterminator activity"

### SUPPLEMENTARY INFORMATION

**A** TETRISv1 *Xist*-2kb random-primed RT

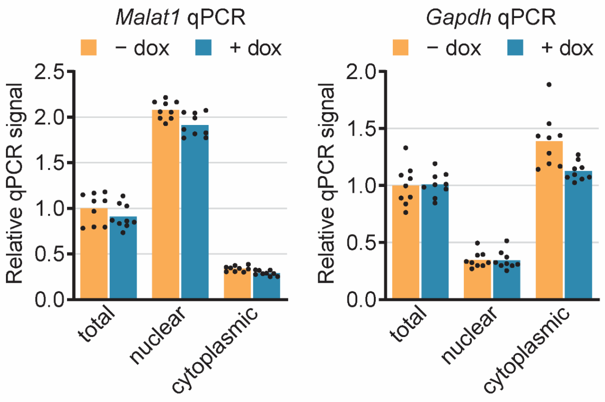


**B** Sanger sequencing of TETRIS plasmids

TETRISv1 *Hottip*

AGCTGTTCCGAGCTTGTGAGGGAGCCTCCAGCTGCAGTCCTTTGTCTGTGGGACTAGATCCCTTCATGGCCCAATGCTCTGGGAAACGAAG
CAGCTCAATACAAACTTTTGATATTAGTTTTTTAAAGGTATTTACACTTTGGTGAGAGTAGCTGGGTACGGAAGCTGCTTCGTTAATCAGC
CCCCGACCCCGACCTTAGGTTCGGCTGGGACTAGGAGCTGGCCGGAAAAGAAAAAAAGCTTCGGGAAGTCGGAGAGAATGAAAAACAAAAC
AAAACTGCACAAGGTCATTTTCTGTAAATCGGATCCGCGGCCGCGATATCGCTAGCTCGAGAATTCGACTGTGCCTTATCGATAAGCTTGT
CGACGATATCTCTAGAGGATCATAATCAGCCATACCACATTGTAGAGGTCTTACTTGCTTTAAAAAACCTCCCACACCTCCCCCTGAACCT
GAAACATAAAATGAATGCAATTGTTGTTGTTAACTTGTTTATTGCAGCTTATAATGGTTACAAATAAAGCAATAGCATCACAAATTTCACA
AATAAAGCATTTTTTTCACTGCCTCGAGCTTCCTCGCTCACTGACTCGCTGCGCTCGGTCGTTCGGCTGCGGCGAGCGGTATCAGCTCACT
CAAAGGCGGTAATACGGTTATCCACAGAATCAGGGGAT

TETRISv1 *Xist*-2kb

TAGTCATCATTTTTTCGAAGTGCCTGCCCAGGTCGGGAGAGCGCATGCTTGCAATTCTAACACTGAAGTGTTGGATGATGTCGGATCCGAT
TCGAGAGACCGAGGCTGCGGGTTCTTGGTCGATGTAAATCATTGAAACCTCACCTATTAAAAGAAAGAAAAGTATCTAAGGCCATTTCAAG
GACATTTGACTCATCCGCTAAATCGGATCCGCGGCCGCGATATCGCTAGCTCGAGAATTCGACTGTGCCTTATCGATAAGCTTGTCGACGA
TATCTCTAGAGGATCATAATCAGCCATACCACATTGTAGAGGTCTTACTTGCTTTAAAAAACCTCCCACACCTCCCCCTGAACCTGAAACA
TAAAATGAATGCAATTGTTGTTGTTAACTTGTTTATTGCAGCTTATAATGGTTACAAATAAAGCAATAGCATCACAAATTTCACAAATAAA
GCATTTTTTTCACTGCCTCGAGCTTCCTCGCTCACTGACTCGCTGCGCTCGGTCGTTCGGCTGCGGCGAGCGGTATCAGCTCACTCAAAGG
CGGTAATACGGTTATCCACAGAATCAGGGGAT

TETRISv2 *Xist*-2kb

GCATGCTTGCAATTCTAACACTGAAGTGTTGGATGATGTCGGATCCGATTCGAGAGACCGAGGCTGCGGGTTCTTGGTCGATGTAAATCAT
TGAAACCTCACCTATTAAAAGAAAGAAAAGTATCTAAGGCCATTTCAAGGACATTTGACTCATCCGCTAAATCGGATCCGCGGCCGCGATA
TCGCTAGCTCGAGAATTCGACTGTGCCTTATCGATAAGCTTGTCGACGATATCTCTAGAGGATCATAATCAGCCATACCACATTGTAGAGG
TCTTACTTGCTTTAAAAAACCTCCCACACCTCCCCCTGAACCTGAAACATAAAATGAATGCAATTGTTGTTGTTAACTTGTTTATTGCAGC
TTATAATGGTTACAAATAAAGCAATAGCATCACAAATTTCACAAATAAAGCATTTTTTTCACTGCCTCGAGCTTCCTCGCTCACTGACTCG
CTGCGCTCGGTCGTTCGGCTGCGGCGAGCGGTATCAGCTCACTCAAAGGCGGTAATACGGTTATCCACAGAATCAGGGGATAACGCAGGAA
AGAACATGTGATGGAAGACAATAGCAGGCATGCTGGGGATGCGGTGGGCTCTATGGCCCGGTTAATTAACTAGAGCTCGCTGATCAGCCT
CGACTGTGCCTTCTAGTTGCCAGCCATCTGTTGTTTGCCCCTCCCCCGTGCCTTCCTTGACCCTGGAAGGTGCCACTCCCACTGTCCTTTC
CTAATAAAATGAGGAAATTGCATCGCATTGTCTGAGTAGGTGTCATTCTATTCTGGGGGGTGGGGTGGGGCAGGACAGCAAGGGGGAGGAT

**Supplementary Figure S1. Fractionation controls and Sanger sequencing of polyadenylation sequences in TETRIS plasmids.** (**A**) To assess subcellular fractionation and evenness of loading, *Malat1* RNA and *Gapdh* mRNA levels were measured via qPCR in the same randomly-primed cDNA samples from whole-cell (total), nuclear, and cytoplasmic RNA used in Figure 1E-F. Dots represent individual technical replicate measurements, and bars represent the average value. (**B**) Sanger sequencing of TETRISv1 *Hottip*, TETRISv1 *Xist*-2kb, and TETRISv2 *Xist*-2kb plasmids. Cyan highlights *Hottip* or *Xist*-2kb cargo sequence, red highlights the SV40 early polyadenylation sequence (same as in pTRE-Tight and other common expression plasmids), and green highlights the BGH polyadenylation sequence (1). See Supplementary Table S1 for information regarding experimental replicates and Supplementary Table S2 for oligo sequences.

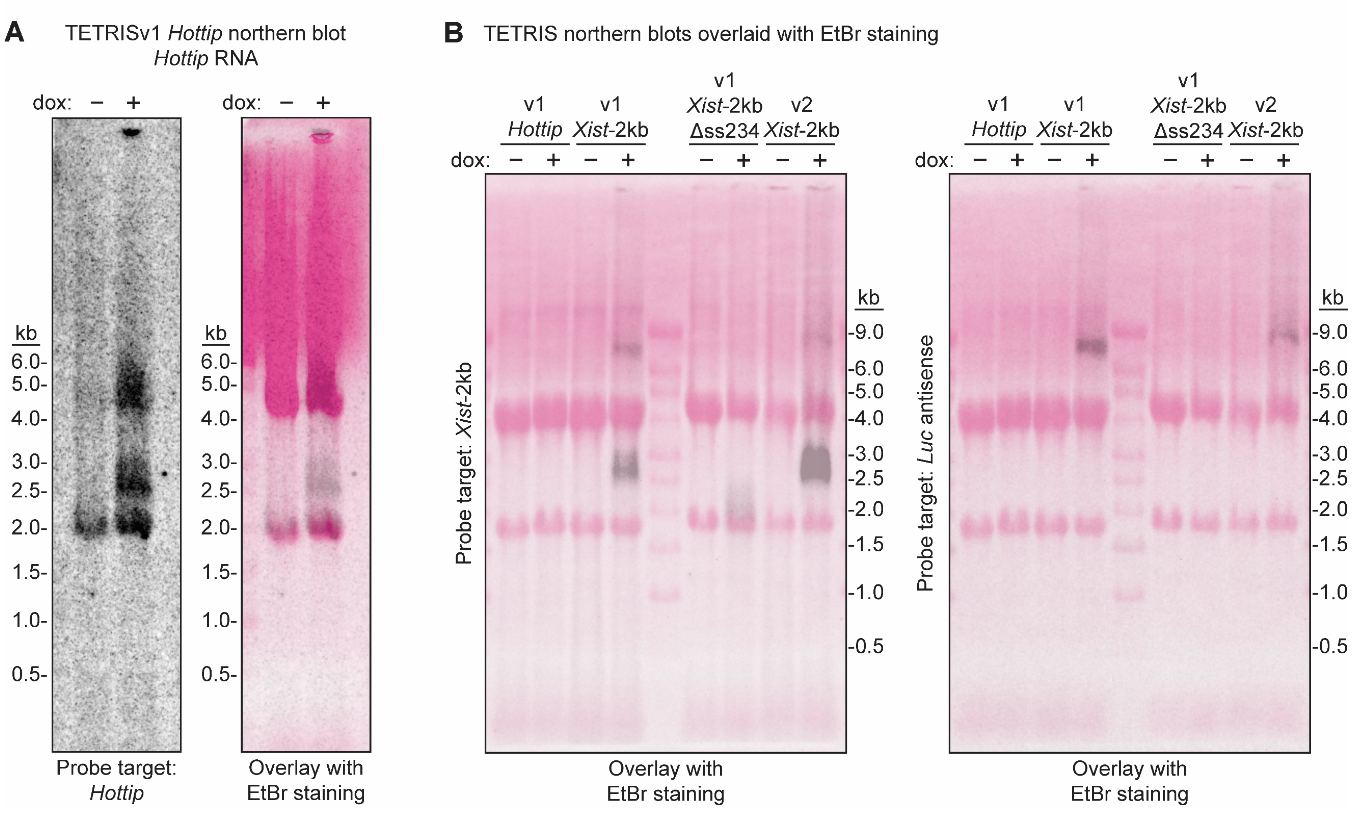


**Supplementary Figure S2. Additional northern blot images related to Figure 2E and F.** (**A**) The same TETRISv1 *Hottip* RNA samples used in Figures 2E-F were analyzed by northern blot using a probe targeting *Hottip* RNA. The left panel shows phosphor image alone, and the right panel shows this phosphor image (black) laid over a fluorogram of the nylon membrane after transfer of ethidium-bromide-stained RNA (magenta), where 18S and 28S rRNAs and Millennium RNA size markers are visible. The two doxycycline-dependent bands correspond with the expected sizes of spliced and unspliced *Hottip* RNA produced from the TETRIS expression cassette (2.2 and 4.2 kb respectively, plus poly(A) tail length). The TETRISv1 *Hottip* transgene has mm10 genomic coordinates chr6:52263006-52266839. (**B**) The phosphor images depicted in Figure 2E-F (black) are laid over ethidium bromide fluorograms (magenta) to show rRNAs and Millennium RNA size markers.

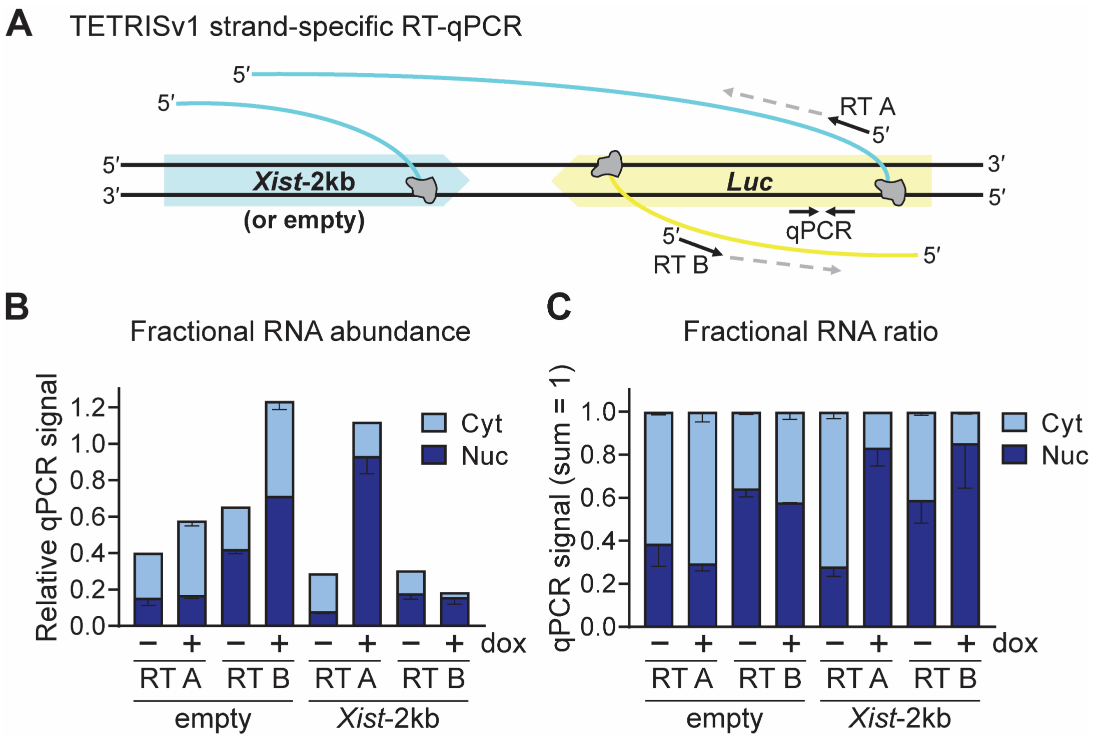


**Supplementary Figure S3. In TETRISv1, *Xist*-2kb induction causes an increase in *Luc* antisense transcripts in the nucleus and a decrease of *Luc* mRNAs in the cytoplasm.** (**A**) Diagram depicting individual strands of TETRISv1 DNA (black) being transcribed (grey Pol II complexes) to produce different RNAs (cyan and yellow). Positions of strand-specific reverse transcription primers and qPCR primers over the *Luc* gene are shown as black arrows. (**B**) Cells harboring TETRISv1 empty (no cargo) or *Xist*-2kb were treated with or without doxycycline, and RNA was prepared from cytoplasmic and nuclear fractions. Relative qPCR signal is shown from the strand-specific RT-qPCR strategy described in (A). (**C**) Sums of nuclear and cytoplasmic qPCR signal for each cell identity in (B) were normalized to 1 to show changes in the nuclear ratio of signal. See Supplementary Table S1 for information regarding experimental replicates and Supplementary Table S2 for oligo sequences.

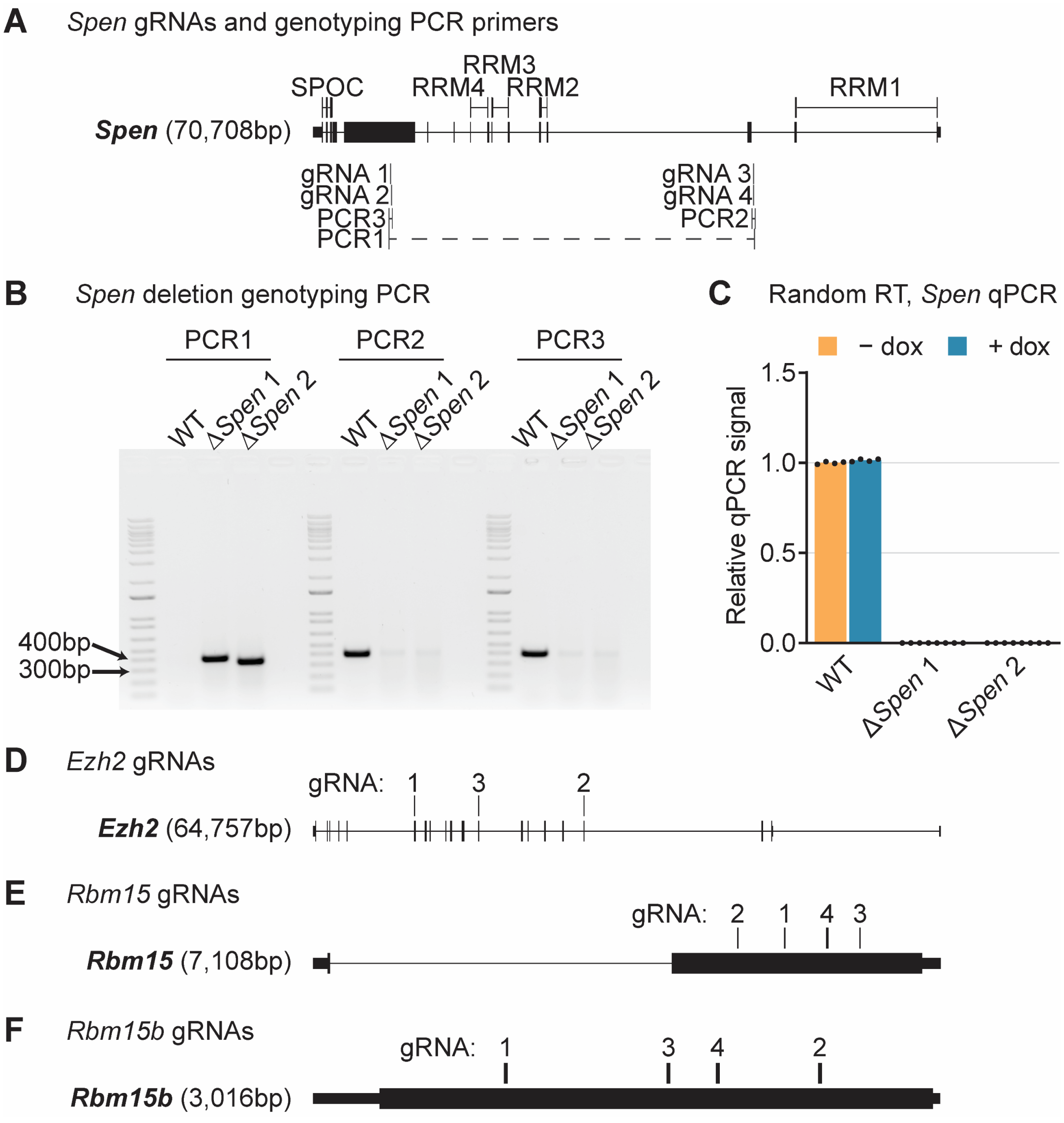


**Supplementary Figure S4. *Spen*, *Rbm15*, *Rbm15b*, and *Ezh2* CRISPR targeting strategies.** (**A**) Gene diagram showing the location of *Spen* exons, annotated functional domains (above the gene), sgRNA target locations (below the gene), and genotyping PCR products (below the sgRNA locations). (**B**) *Spen* deletion genotyping PCR showing the presence of deletion-specific bands and absence of wild-type-specific bands in genomic DNA from two independent homozygous deletion clones. (**C**) RT-qPCR confirming absence of *Spen* mRNA in deletion clones. Dots represent individual technical replicate measurements, and bars represent the average value. (**D-F**) Gene diagrams showing the locations of exons and sgRNA target locations for *Rbm15* (D), *Rbm15b* (E), and *Ezh2* (F). See Supplementary Table S1 for information regarding experimental replicates and Supplementary Table S2 for oligo sequences.

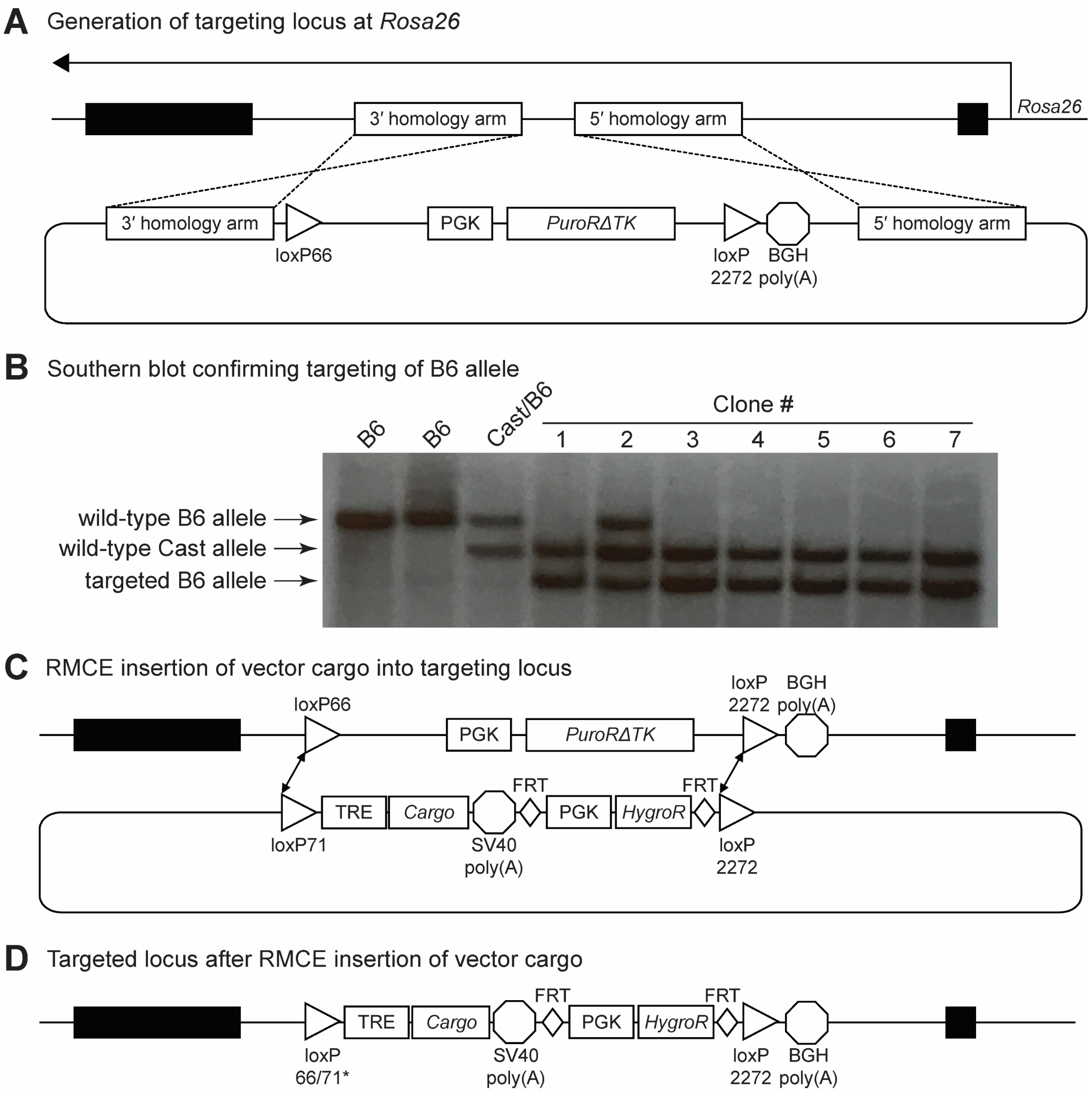


**Supplementary Figure S5. *Rosa26*-RMCE system for inducible expression of inserted transgenes.** (**A**) Generation of targeting locus at *Rosa26*. PGK, constitutive promoter; loxP66 and loxP2272, Cre recombinase recognition sequences; *PuroRΔTK*, gene encoding puromycin resistance (for positive selection) and truncated thymidine kinase (for negative selection with ganciclovir). (**B**) Southern blot confirming successful targeting of the B6 allele, using a probe that detects a region 3′ of the 3′ *Rosa26* homology arm. “Clone #1” was expanded and used for the RMCE experiments in this work. (**C**) RMCE insertion of pCARGO vector cargo into targeting locus at *Rosa26*. loxP71, Cre recombinase recognition sequence for recombination with loxP66. TRE, tetracycline responsive element (the doxycycline inducible promoter). *HygroR*, gene encoding resistance to hygromycin. FRT, Flp recombinase target. (**D**) Targeted locus after RMCE insertion of cargo. loxP66/71*, recombined site no longer usable by Cre recombinase, preventing removal of cassette.

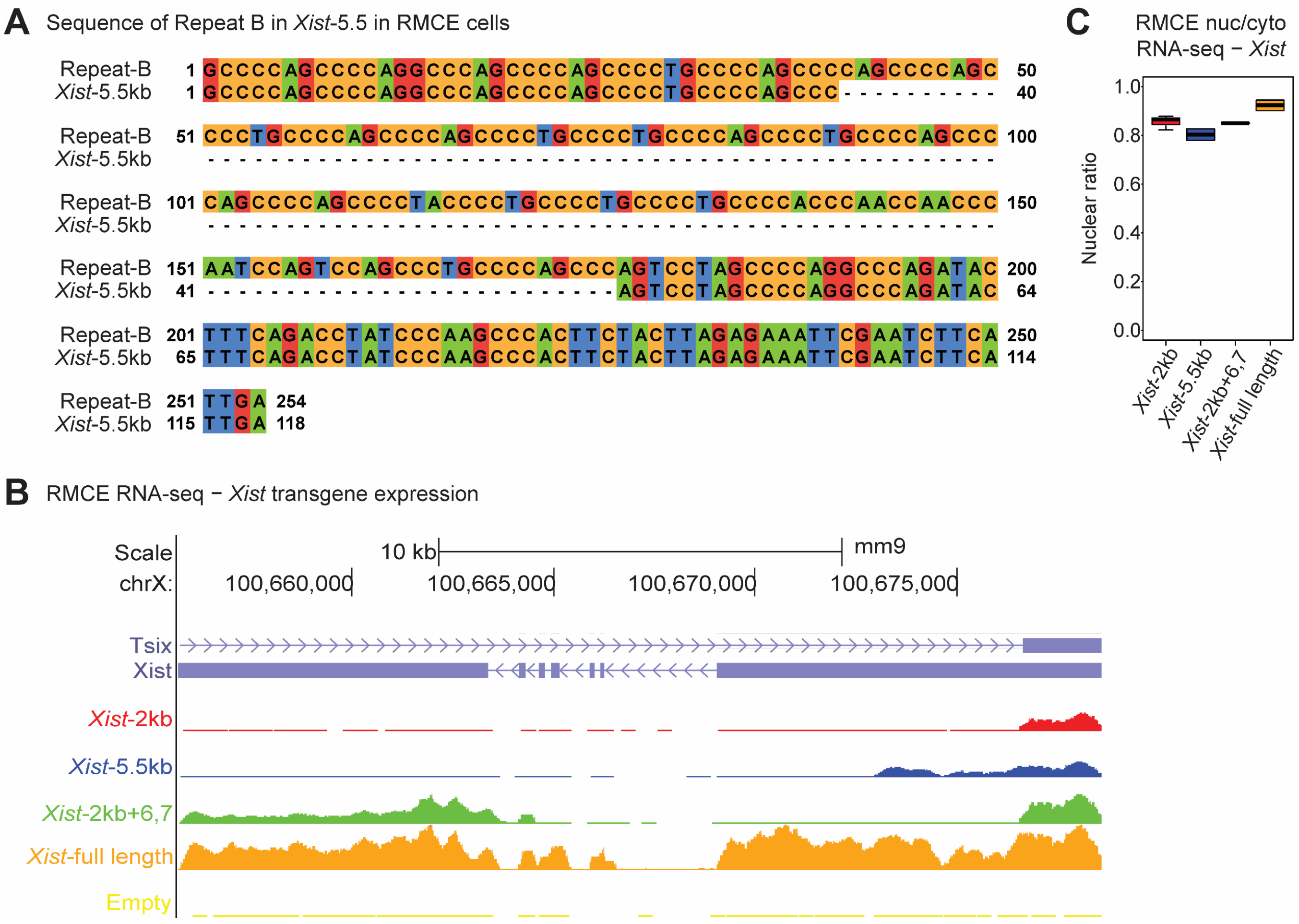


**Supplementary Figure S6. *Xist* transgenes are expressed from the *Rosa26* locus.** (**A**) The *Xist*-5.5kb transgene aligned to the Repeat-B portion of endogenous *Xist* (254bp total) shows a deletion of 136bp. (**B**) Wiggle tracks showing nuclear *Xist* read density in one representative replicate of each *Xist* transgene line. Reads are aligned to the endogenous *Xist* locus, which is not expressed in these cells (as seen in the empty-cargo line). (**C**) Nuclear ratio of reads mapping to *Xist* in each line expressing *Xist* transgenes.

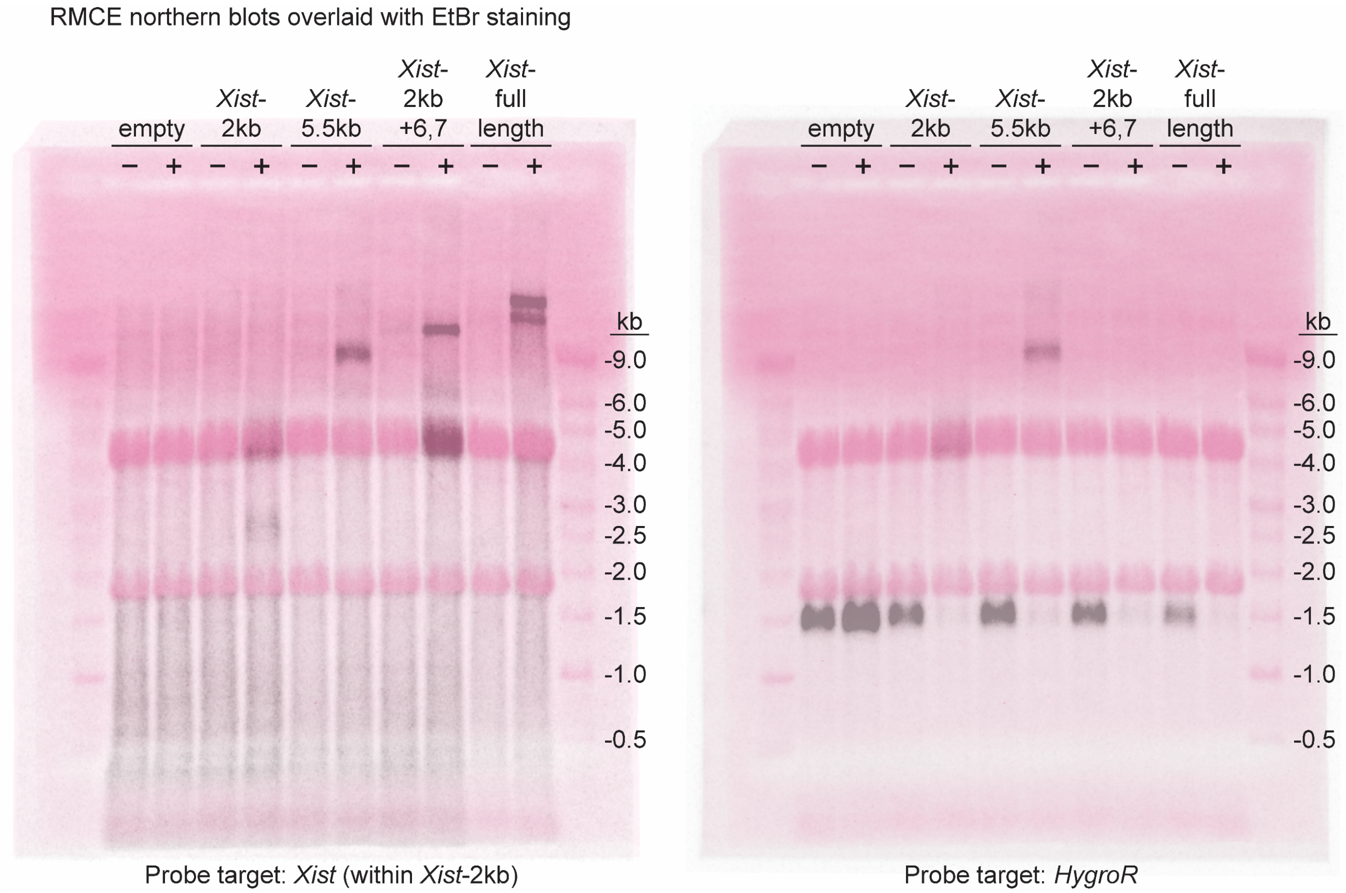


**Supplementary Figure S7. Additional northern blot images related to Figure 5H and I.** The phosphor images depicted in Figure 5H-I (black) are laid over ethidium bromide fluorograms (magenta) to show rRNAs and Millennium RNA size markers.

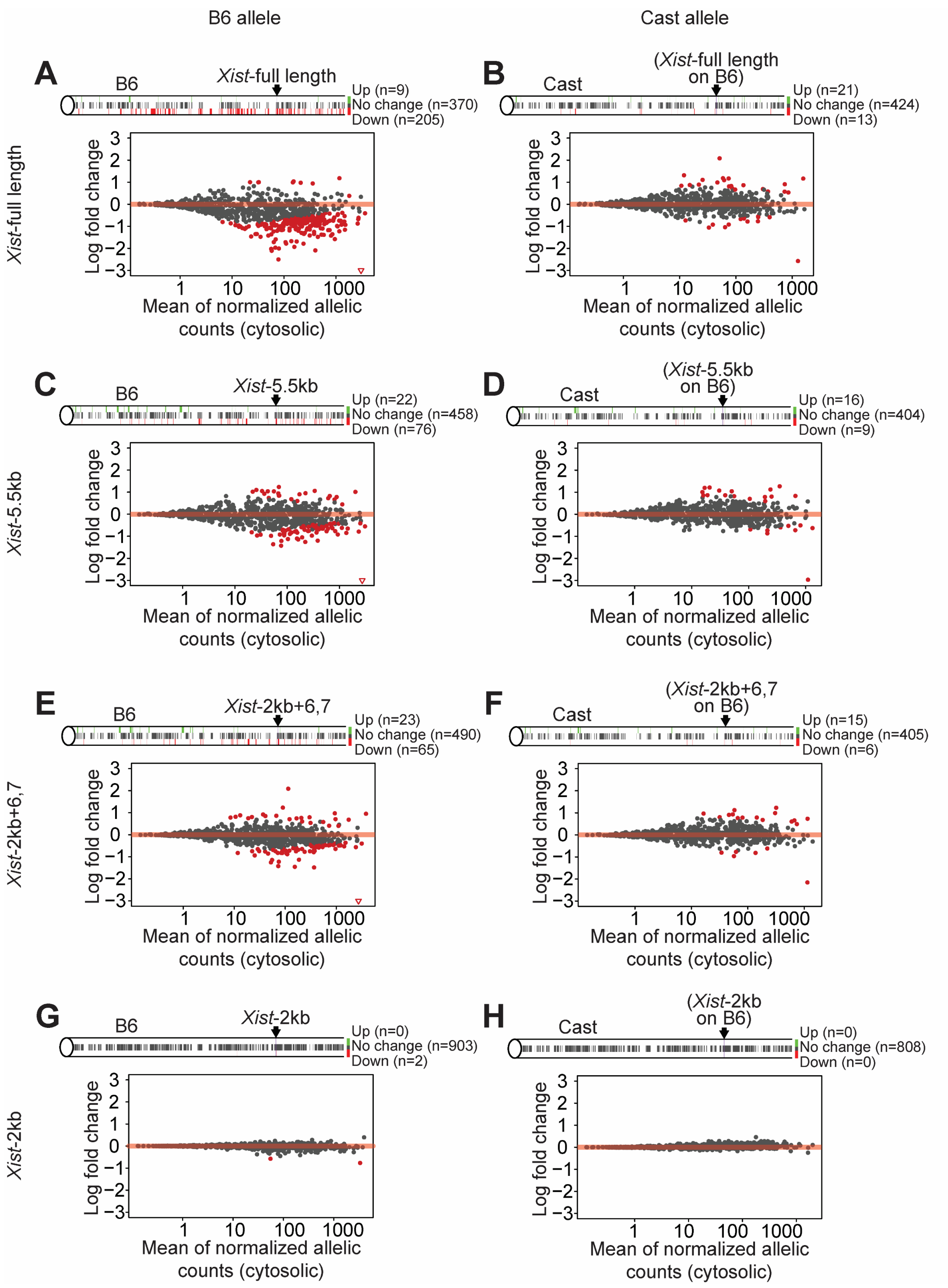


**Supplementary Figure S8. Locations and levels of genes silenced by *Xist* transgenes.** (**A**) MA plot showing differential expression between empty-cargo cells and full-length *Xist* cells on the B6 allele of chromosome 6. Log fold change was plotted against the mean of normalized allelic cytoplasmic counts. Each dot is an expressed gene, and red dots show significantly changing genes as determined by DESeq2 analysis (adjusted p value < 0.05). Red triangle indicates gene below the lower y-axis limit. Locations of all expressed genes along chromosome 6 are shown above the plot. Genes which increase, decrease, or do not change in full-length *Xist* cells are marked by green, red, and grey bars, respectively. The location of *Xist* on the B6 allele is indicated. (**B**) Similar to (A) but for the Cast allele. For comparison, the transgene location from the B6 allele is indicated. (**C-H**) Similar to (A-B) but for expression of *Xist*-5.5kb (C-D), *Xist*-2kb+6,7 (E-F), and *Xist*-2kb (G-H). See Supplementary Table S5 for RNA-seq gene expression data.

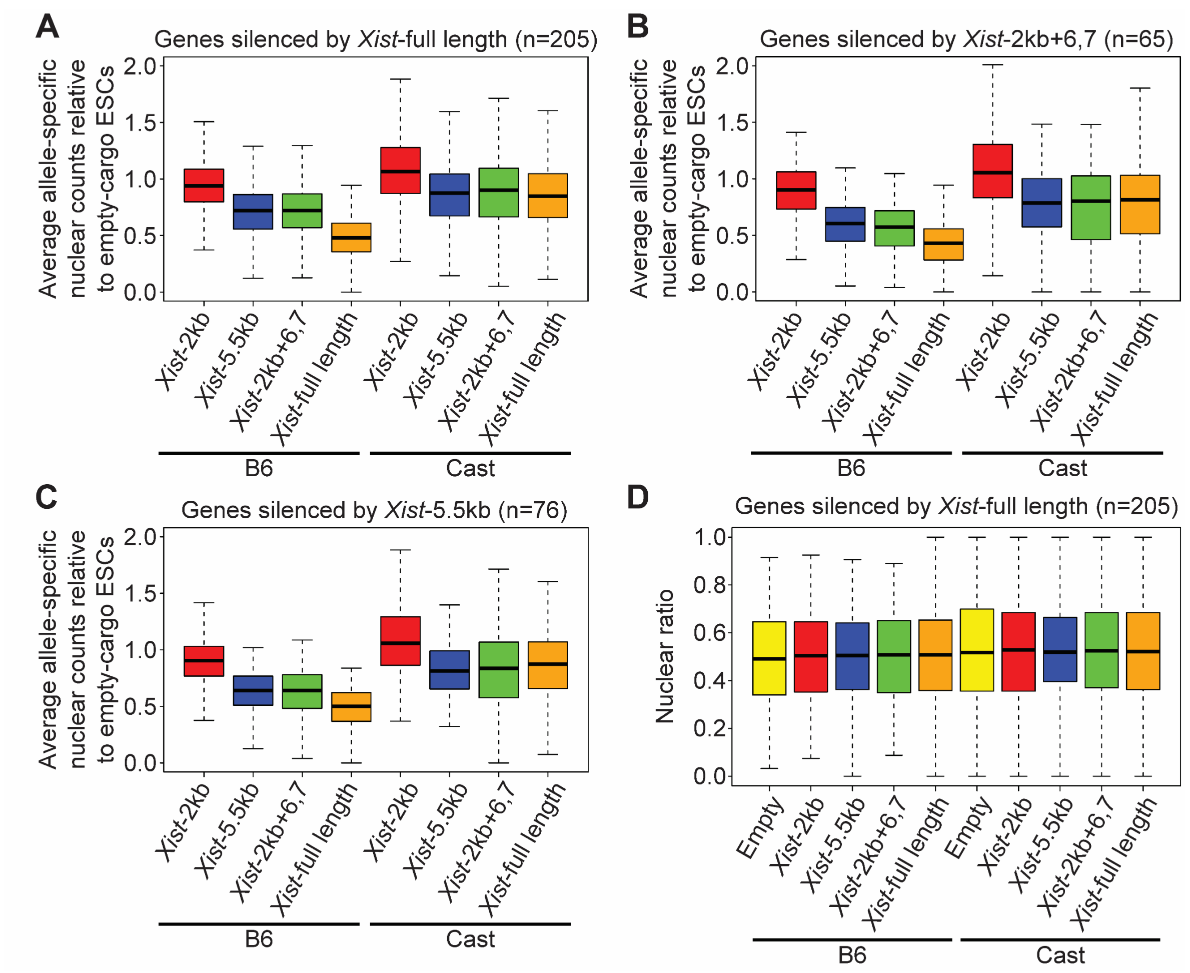


**Supplementary Figure S9. *Xist* transgenes silence genes across chromosome 6 at the transcriptional level.** (**A-C**) Average allele-specific normalized nuclear counts relative to empty-cargo ESCs for the 205 genes significantly repressed by full-length *Xist* (A), the 65 genes significantly repressed by *Xist*-2kb+6,7 (B), or the 76 genes significantly repressed by *Xist*-5.5kb (C). (**D**) Average nuclear ratio of allele-specific reads for the 205 genes significantly repressed by full-length *Xist*. See Supplementary Table S5 for RNA-seq gene expression data.

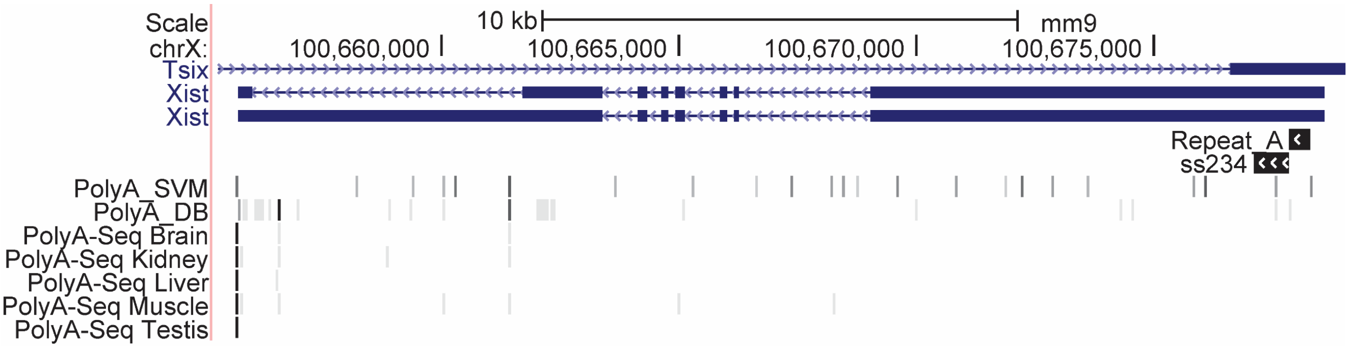


**Supplementary Figure S10. Polyadenylation sites are predicted throughout *Xist*, but polyadenylation is detected primarily at its 3′ end.** UCSC Genome Browser view of mouse *Xist* (on the negative strand, with its 5′ end on the right), showing locations of polyadenylation sites predicted by PolyA_SVM (2). Darker shades of grey correspond to sites that are predicted to be stronger (i.e., lower PolyA_SVM score (2)). Positions of experimentally observed polyadenylation sites are shown below, cataloged by PolyA_DB (3) and by the PolyA-Seq tracks available in the UCSC Genome Browser (Adnan Derti and Tomas Babak, Merck Research Laboratories, GSE30198; (4)). For these tracks, darker shades of grey correspond to more intense read signals.

**Supplementary Table S1. Experimental replicates.** Table contains information regarding numbers of biological and technical replicates for each experiment.

**Supplementary Table S2. Sequences and usage of all DNA oligos.** Table contains the sequences of all single-stranded DNA oligonucleotides used in this study as well as their usage in each experiment.

**Supplementary Table S3. CHART-MS – proteins enriched with *Xist*-2kb.** Table contains the 22 proteins identified my mass spectrometry as being enriched with *Xist*-2kb. These proteins fit two criteria: they had spectral counts detected in both *Xist*-2kb replicates, and they had no spectral counts detected in the *Xist*-2kb ΔrA234 negative control.

**Supplementary Table S4. CHART-MS – all proteins.** Table contains total spectrum count data for all proteins identified by CHART-MS in all samples. Proteins are sorted by a *Xist*-2kb pseudo-enrichment score, calculated as the sum of *Xist*-2kb spectral counts divided by the sum of *Xist*-2kb ΔrA234 spectral counts and 0.1 (to prevent division by zero).

**Supplementary Table S5. RMCE RNA-seq data.** Table contains allele-specific raw and normalized nuclear and cytoplasmic counts for all genes detected as differentially expressed in any of the *Xist* transgene lines. Sheet 1 ('all_changers') contains raw allele-specific read counts for each replicate of each cell line for every gene detected as significantly differentially expressed in any of the *Xist* transgene lines compared to the empty line. Sheet 2 ('all_changers_norm') contains the upper quartile normalized counts for each gene and line as in Sheet 1. Sheet 3 ('deseq2_results') contains the log 2-fold changes ('L2FC') and adjusted p-values ('PAdj') for each DESeq2 comparison for each gene detected as significantly differentially expressed in any of the *Xist* transgene lines.

**Supplementary File 1. CHART-MS data.** File (.sf3 format) for viewing in Scaffold. Contains all compiled CHART-MS data.

### Supplementary Methods

#### *Embryonic stem cell culture*

E14 ESCs (kind gift of D. Ciavatta) were grown on gelatin coated dishes at 37°C in a humidified incubator at 5% $CO_2$. Medium was changed daily and consisted of DMEM high glucose plus sodium pyruvate, 15% ESC qualified fetal bovine serum, 0.1 mM non-essential amino acids, 100 U/mL penicillin-streptomycin, 2 mM L-glutamine, 0.1 mM 2-mercaptoethanol, and 1:500 LIF conditioned media produced from Lif-1Cα (COS) cells (kind gift of N. Hathaway). ESCs were split at an approximate ratio of 1:6 every 48 h. *Rosa26*-RMCE cells were grown on gamma-irradiated mouse embryonic fibroblast (MEF) feeder cells plated at approximately 1.5 million cells per 10-cm plate. Prior to harvesting of RNA for sequencing, *Rosa26*-RMCE cells were passaged twice off of MEF feeder cells with a 40-min pre-plate each passage and grown in 70% MEF-conditioned medium supplemented as above.

#### *TETRIS line generation*

TETRIS lines were made as described in (5). Briefly, 500,000 E14 cells were seeded in a single well of a 6-well plate and transfected 24 h later with 0.5 µg pTETRIS plasmid, 0.5 µg *rtTA*-cargo, and 1 µg of pUC19-piggyBac transposase using Lipofectamine 3000 (Invitrogen) according to manufacturer instructions. Cells were selected for 7-9 days with puromycin (2 µg/mL) and G418 (200 µg/mL) beginning 24 h after transfection. TETRIS cargo vectors were generated as part of (5).

#### *TETRIS luminescence assays*

For each independent TETRIS cell line, six wells of a 24-well plate were seeded at 100,000 cells per well. Three of the six wells were induced with 1 µg/mL doxycycline (Sigma) beginning when the cells were plated, and the remaining three wells served as “no dox” controls. After 48 h, the cells were washed with PBS and lysed with 100 µL of passive lysis buffer (Promega) and luciferase activity was measured using Bright-Glo™ Luciferase Assay reagents (Promega) on a PHERAstar FS plate reader (BMG Labtech). Luciferase activity was normalized to total protein concentration in the lysates measured via Bradford assay (Bio-Rad).

#### *RNA isolation and subcellular fractionation*

Cells were plated on 6-well or 12-well plates at an approximate density of 400,000 or 200,000 cells per well, respectively. Cells were induced with doxycycline at 1 µg/mL for 2 d unless otherwise stated. All steps in RNA isolation were performed at 4 °C. For whole-cell (total) RNA samples, cells were washed twice with 1 mL ice-cold PBS, and RNA was isolated directly from cells using Trizol according to manufacturer protocol (Invitrogen). To isolate RNA from cytoplasmic and nuclear fractions, cells were washed twice with 1 mL ice-cold PBS, scraped in 1 mL ice-cold PBS, centrifuged at 1500 x g for 5 min, and resuspended in 250 µL low salt solution (10 mM KCl, 1.5 mM $MgCl_2$, 20 mM Tris-HCl pH 7.5) supplemented with 1 mM PMSF, 1 mM DTT, and 1:100 protease inhibitor cocktail (PIC; Sigma P8340). Triton X-100 was added to a final concentration of 0.1%, cells were rotated for 10 min, and tubes were then centrifuged for 5 min at 1500 x g. 200 µL of supernatant was added to 1 mL Trizol (cytoplasmic fraction). The remaining supernatant was discarded and the nuclear pellet was washed by rotating for 2 min in low salt solution without Triton X-100 and centrifuged at 1300 rpm (Eppendorf 5415D) for 10 min. Nuclear pellets, which at this stage contained both soluble nuclear and chromatin-bound fractions, were resuspended in 1 mL Trizol. Isolation of RNA from Trizol was performed according to manufacturer protocol (Invitrogen).

TRIzol samples were stored at -80 °C before thawing on the day of RNA preparation. To prepare RNA, 200 µL chloroform was added to all TRIzol samples, which were then shaken vigorously for 15 s and then incubated at room temperature for 3 min before centrifuging at 16,100 x g for 20 min at 4 °C. The top aqueous phase (420 µL) was added to 500 µL ice-cold 100% isopropanol and mixed well via vortex before pelleting precipitated RNA at 16,100 x g for 20 min at 4 °C. RNA pellets were washed once with ice-cold 70% (v/v) ethanol and centrifuged 16,100 x g for 20 min at 4 °C. After complete removal of ethanol supernatant (via pipet and ~10-15 min of air-drying), ice-cold RNase-free water was added to RNA pellets and incubated on ice for at least two hours before thoroughly resuspending via pipet. RNA was quantified using a NanoDrop Lite spectrophotometer (Thermo Scientific). RNA samples were periodically run on an agarose gel to assess RNA integrity.

***RT-qPCR***

For random-primed reverse transcription, equal amounts of RNA (typically 0.5-1 µg) were reverse transcribed in 10 or 20 µL reactions using the High-Capacity cDNA Reverse Transcription Kit (Applied Biosystems) according to manufacturer protocol with random primers. For strand-specific reverse transcription in Supplementary Figure S3B and C, 100 ng RNA was used with 1 µM strand-specific primer (omitting the random primers). Products of reverse transcription reactions were diluted 2- to 10-fold with water, of which 1 µL was used as cDNA template for 10-µL quantitative PCR reactions containing 0.5 µM each forward and reverse primer and 1X Bio-Rad iTaq Universal SYBR Green Supermix. Specificity of primer pairs was confirmed using the *in silico* PCR tool on the UCSC Genome Browser, by analyzing melt curves of qPCR products, and by running control samples expected to not contain the target sequence. Quantitative PCR was performed on a Bio-Rad C1000 Touch Thermal Cycler equipped with a Bio-Rad CFX96 Real-Time System with the following thermocycling parameters: initial denaturation at 95 °C for 10 min; 40 cycles of 95 °C for 15 sec, 60 °C for 30 sec, and 72 °C for 30 sec followed by a plate read. Control reactions without RT enzyme were periodically performed to assess DNA contamination. Relative quantities of RNA (qPCR signal) were determined using Bio-Rad CFX Maestro software using "regression" Cq determination mode and by fitting samples to a simultaneously-run serial dilution curve to accurately determine qPCR efficiency. Data were normalized (usually to the average of the corresponding “no dox” control measurements) using Microsoft Excel and plotted using GraphPad Prism 8. See Supplementary Table S1 for information regarding experimental replicates.

***Puromycin sensitivity assay***

TETRIS cells expressing *Xist*-2kb were grown for 2 days with or without doxycycline (1 μg/mL) before equal numbers of cells (500,000) were plated on 6-well plates in medium containing puromycin (2 μg/mL) with or without doxycycline. Cells were trypsinized, quenched with culture medium, mixed with trypan blue dye, and counted after 24 h and 48 h of puromycin treatment. Cell survival is shown relative to cells grown without doxycycline.

***Stellaris single-molecule sensitivity RNA FISH***

Custom Stellaris FISH probes were designed against the first 2 kb of *Xist*, firefly luciferase (*luc2* in pGL4.10; Promega), and the hygromycin resistance gene (*HygroR*) using the Stellaris RNA FISH Probe Designer (Biosearch Technologies, Inc.) and labeled with Quasar 670 (*Xist*-2kb) or 570 (*Luc* and *HygroR*) dye. Cells were grown on glass coverslips for 2 days in the presence or absence of 1 μg/mL doxycycline before being washed once with 1X PBS, fixed for 10 min at room temperature with 4% formaldehyde in 1X PBS, washed twice with 1X PBS, and permeabilized overnight with cold 75% ethanol at 4 °C. 1 μL of 2.5 μM (*Xist*-2kb) or 25 μM (luciferase and *HygroR*) probe was added to 100 μL of hybridization solution (10% dextran sulfate, 2X SSC, 10% formamide) and pre-warmed to 37 °C. Coverslips were washed at 37 °C for 2-5 min in pre-warmed wash buffer (2X SSC, 10% formamide). Coverslips were incubated with diluted probes overnight at 37 °C in a humidified chamber, then washed twice with wash buffer at 37 °C for 30 min, adding DAPI to 5 ng/mL for the second wash. Coverslips were rinsed with 2X SSC, mounted using Prolong Gold and allowed to cure overnight at room temperature or using Vectashield with nail polish sealing the edges.

Multi-channel Z-stack images were captured on an Olympus BX61 widefield fluorescence microscope using a Plan-Aprochromat 63X/1.4 oil objective and a Hamamatsu ORCA R2 camera, controlled by Volocity 6.3 software. Excitation was provided by a mercury lamp and the following filters were used for the three fluorescent channels that were imaged: 377/25 ex, 447/30 em for DAPI (DAPI-5060B Semrock filter); 562/20 ex, 642/20 em for Quasar 570 (Semrock TXRED-4040B filter); 628/20 ex, 692/20 em for Quasar 670 (Semrock Cy5 4040A filter). Pixel size was 0.108 μm, Z spacing was 0.2 μm, and images had 1344x1024 pixels. Approximately 40 Z-stacks were acquired for each image. Z-stacks were deconvolved using the iterative-constrained algorithm (Mediacy AutoQuantX3) with default algorithm settings. Sample settings for the deconvolution were: peak emissions for dyes (670 nm, 565 nm, 461 nm for Quasar 670, Quasar 570, and DAPI respectively), widefield microscopy mode, NA = 1.4, RI of oil = 1.518, and RI of sample = 1.45. Images are shown as maximum intensity projections made using ImageJ.

***Generation of TETRISv2 plasmids***

A sequence comprised of the EF1α promoter, the puromycin resistance gene, and an SV40 early poly(A) site was synthesized by GeneArt (Thermo Fisher) and cloned into the SgfI and NsiI sites in pTETRISv1, generating pTETRISv1.5. This sequence replaced the EF1α-*CymR*-*PuroR*-SV40earlypoly(A) sequence of TETRISv1, effectively removing the *CymR* sequence. Subsequently, a sequence was synthesized by GeneArt that contained an SV40 early poly(A) site followed by a BGH poly(A) site, then followed by a 1.1 kb region downstream and proximal to the natural termination sequence of the mouse *Pgk1* gene (corresponding to chrX:103,399,675-103,400,826 on the UCSC mm9 genome build). This sequence was then cloned into the ClaI and BsrG1 sites of pTETRISv1.5 to generate pTETRISv2. *Xist*-2kb was then cloned into the SwaI site of pTETRISv2 via Gibson assembly as previously described. Plasmid sequences were confirmed via Sanger sequencing.

***Generation of TETRISv3 plasmids***

pTETRISv2 plasmid was digested with NEB XhoI and PmeI to remove the lncRNA cargo sequence (including the TRE promoter and both SV40 early and BGH poly(A) signals) from the plasmid backbone. The plasmid backbones were blunted with the NEB Quick Blunting Kit and re-ligated with NEB Quick Ligase. Removal of the lncRNA cargo sequence was confirmed via Sanger sequencing. Using TETRISv1 plasmids as templates, TRE-*Xist*-2kb-SV40earlypoly(A) and TRE-empty-SV40earlypoly(A) sequences were PCR-amplified with primers (DL673 and DL675) containing BstXI sites. These PCR products were digested with NEB BstXI and inserted into lncRNA-cargo-removed TETRISv2 backbone that had been digested with NEB BstXI and desphosphorylated with NEB Antarctic Phosphatase. This ligation step was performed using NEB Quick Ligase. Plasmid sequences were confirmed via Sanger sequencing.

***Northern blots***

Northern blots were performed using a protocol adapted from (6). Using a Thermo Fisher Owl EasyCast B2 gel system, a 150-mL gel was prepared containing 1.5 g agarose (Genesee Scientific 20-102QD; to 1.0% w/v), 135 mL RNase-free water, 15 mL 10X NorthernMax Denaturing Gel Buffer (Thermo Fisher AM8676), and 10 µL 1% (w/v) ethidium bromide (Fisher BP1302-10). The gel was poured in a chemical fume hood and allowed to rest for at least 1 h. For each sample to be analyzed, 15 µg RNA was brought up to 10 µL with RNase-free water and then mixed with 25 µL Formaldehyde Load Dye (Thermo Fisher AM8552). In parallel, 10 µL Millennium RNA Markers (Thermo Fisher AM7150) was mixed with 25 µL Formaldehyde Load Dye. Samples were incubated at 65 °C for 15 min and then placed on ice for at least 5 min. Prior to loading samples into the gel, the gel was pre-run at 93 V for 15 min in 1X NorthernMax Running Buffer (Thermo Fisher AM8671). In the chemical fume hood, the gel was loaded with samples, covered with aluminum foil to protect from light, and electrophoresed at 93 V in the same buffer until the bromophenol blue dye front was ~2 cm from the end of the gel (~3.5 h). To check quality of electrophoresis, the gel was imaged with a Bio-Rad ChemiDoc MP Imaging System in ethidium bromide mode (trans UV). The gel was washed twice with ~250 mL RNase-free water and then twice with ~250 mL 10X SSC (1.50 M NaCl, 150 mM sodium citrate, pH 7.0). Each wash was performed for 15 min on an orbital shaker at room temp. RNA was transferred to nylon membrane overnight (>16 h) using a capillary transfer apparatus assembled on a Thermo Fisher Owl EasyCast B2 gel tank with 10X SSC as transfer buffer; from bottom to top, the following were wet with 10X SSC, stacked, and rolled to remove air bubbles: upside-down gel-casting tray, 29 cm x 14 cm wick of Whatman 3MM chromatography paper (GE Healthcare 3030-392), upside-down RNA gel (approx. 12 cm x 14 cm), 12.5 cm x 14.5 cm Hybond-N membrane (GE Healthcare RPN303N), three 12 cm x 14 cm sheets of Whatman 3MM chromatography paper, and a ~6-cm-high stack of paper towels cut to 12 cm x 14 cm. As a weight, the lid of a small styrofoam box and an unopened box of P1000 pipet tips were placed on top.

Following transfer, the membrane was placed RNA-side-up in a Stratagene UV Stratalinker 1800 and UV-crosslinked with the auto-crosslink setting (1200 x 100 µJ). To visualize ethidium-bromide-stained RNA, the membrane was imaged with a Bio-Rad ChemiDoc MP Imaging System in Flamingo mode (blue epi, 590/100 nm filter). The membrane was rolled and placed in a hybridization tube with the RNA side facing inward. To the tube, 20 mL ULTRAhyb-Oligo (Thermo Fisher AM8663; pre-warmed to 50 °C and mixed well) was added, and the tube was rotated at 42 °C in a hybridization oven for at least 45 min to pre-hybridize. Meanwhile, a single-stranded DNA oligo (Integrated DNA Technologies; see Supplementary Table S2 for sequences) was labeled with phosphorus-32 ($^{32}$P) in a 20-µL reaction containing 11.5 µL RNase-free water, 1.5 µL 10 µM DNA oligo, 2 µL 10X T4 PNK Buffer, 4 µL 3000Ci/mmol 10mCi/mL [γ-$^{32}$P]ATP (PerkinElmer BLU502A250UC), and 1 µL T4 polynucleotide kinase (New England Biolabs B0201S). The reaction was incubated at 37 °C for ~1 h, 30 µL RNase-free water was added, and labeled DNA probe was purified using a MicroSpin G-50 column (MilliporeSigma GE27-5330-01). To the hybridization tube containing the membrane and 20 mL ULTRAhyb-Oligo, 30 µL labeled probe was added, and the tube was rotated at 42 °C in a hybridization oven overnight (typically >16 h). The membrane was washed twice with 35 mL Northern Wash Buffer (2X SSC, 0.5% sodium dodecyl sulfate (w/v); pre-warmed to 42 °C), 30 min per wash, rotating at 42 °C. The membrane was wrapped in plastic wrap and exposed to a GE Healthcare storage phosphor screen (Kodak SO230) for ~24 h. The storage phosphor screen was imaged with an Amersham Typhoon 5 Biomolecular Imager (GE Healthcare 29187191) in Phosphor Imaging mode with 4000 sensitivity and 100 µm pixel size. Brightness and contrast of images were linearly adjusted (if necessary) in Bio-Rad Image Lab 6.0.1. To estimate molecular weights of RNAs detected by phosphor imaging, images were overlaid with ethidium-bromide-stained (Flamingo mode) images in Adobe Illustrator by adjusting image opacity, allowing co-visualization of Millennium RNA size markers. To strip the membrane for re-probing, 1 L RNase-free water was heated to boiling, and 10 mL 10% (w/v) sodium dodecyl sulfate was added and mixed well. The membrane was incubated for at least 15 min in this solution with occasional stirring before repeating once more. After stripping, the membrane was wrapped in plastic wrap, exposed to storage phosphor screen for ~24 h, and phosphor-imaged again to ensure successful removal of probe signal. Afterward, the membrane was again pre-hybridized in ULTRAhyb-Oligo and probed as described above. Membranes were probed first to detect *Luc* antisense or *HygroR* RNA, and *Xist* RNA was probed after membrane stripping.

#### *SPEN deletion*

To delete SPEN in E14 ESCs, 4 different sgRNAs flanking (2 on each side) an approximately 40-kb genomic region including 3 of the 4 annotated RNA recognition motifs in SPEN were cloned into pX330-U6-Chimeric_BB-CBh-hSpCas9 (a gift from F. Zhang; Addgene plasmid #42230; (7); see diagram in Supplementary Figure S4A and oligo sequences used for cloning in Supplementary Table S2). Flanking sgRNAs were designed using CRISPETa (8). A pool of all 4 sgRNAs each inserted into pX330 were prepared using the PureLink HiPure MidiPrep kit (Invitrogen). Transfections were performed using Lipofectamine 3000 (ThermoFisher) according to the manufacturer's protocol. Briefly, 500,000 E14 cells were plated in one well of a 6-well plate the day before transfecting. 800 ng of the pX330 pool and 200 ng of a plasmid containing a puromycin resistance gene were mixed with 2 µL P3000 reagent and Opti-MEM to 125µL. This mixture was added to 7.5 µL Lipofectamine 3000 reagent diluted in 117.5 µL Opti-MEM and incubated for 5 min at room temperature before adding the entire mixture to the cells in fresh ESC medium. After 24 h, medium was changed and puromycin was added to 2 µg/mL to enrich for cells which received the plasmids. Approximately 66 h after transfection, the cells were plated at low density (500-6000 cells per plate) on 10-cm plates seeded with gamma-irradiated MEFs at 1.5 million cells per plate. Cells were grown for 5-7 d (without puromycin) until colonies were ready to be picked. Freezer stocks and genomic DNA were prepared from the colonies as described for *Rosa26*-RMCE clones below.

Genotyping PCR reactions were performed with genomic DNA (gDNA) using Apex Taq DNA Polymerase (Genesee Scientific). A pair of primers which flank the deletion were used to screen for deletion positive clones (PCR1 in Supplementary Figure S4A-B). Two primer pairs, one pair flanking each set of sgRNA cut sites, were used to screen deletion positive clones for homozygous deletion clones (PCR2 and PCR3 in Supplementary Figure S4A-B). Primer sequences are listed in Supplementary Table S2. Two homozygous deletion clones were expanded and used to generate *Xist*-2kb TETRIS lines following RT-qPCR confirmation of SPEN deletion (Supplementary Figure S4C).

#### *CHART mass spectrometry*

CHART was performed essentially as described in (9), but using the elution protocol from (10). E14 ESCs stably expressing TETRISv1 *Xist*-2kb or TETRISv1 *Xist*-2kb ΔrA234 were grown in the presence of 1 µg/mL doxycycline (Sigma) for 48 h prior to trypsinization, quenched with standard media, washed with PBS, and crosslinked in 1% formaldehyde for 10 min at room temperature. Cells were then flash-frozen in liquid nitrogen for later use. 200 million crosslinked cells were thawed, resuspended in Sucrose Buffer (0.3 M Sucrose, 1% Triton X-100, 100 mM potassium acetate, 10 mM HEPES pH 7.5, 0.1 mM EGTA, 0.5 mM spermidine, 0.15 mM spermine, 1:100 protease inhibitor cocktail [Sigma], 1 mM DTT, 1 U/mL Superase-In [Ambion]) at a concentration of 50 million cells per 4 mL. Cells were transferred to an ice-cold dounce homogenizer and dounced 20 times with a tight pestle with a 5-min rest between the first 10 and last 10 dounce strokes. The suspension was then layered on top of Glycerol Buffer (25% Glycerol, 10 mM HEPES pH 7.5, 1 mM EDTA, 0.1 mM EGTA, 100 mM potassium acetate, 0.5 mM spermidine, 0.15 mM spermine, 1:100 protease inhibitor cocktail [Sigma], 1 mM DTT, 1 U/mL Superase-In [Ambion]) and centrifuged at 1000 x g for 15 min at 4 °C. The process from douncing to centrifugation was repeated once before proceeding with a second round of fixation. After douncing, a pellet of 200 million cells was washed twice with PBS containing 0.5% Tween-20 and then three times with 5 mL Sonication Buffer (50 mM HEPES pH 7.5, 75 mM NaCl, 0.5% N-lauroylsarcosine, 0.1% sodium deoxycholate, 0.1 mM EGTA, 1:100 protease inhibitor cocktail [Sigma], 1 mM DTT, 0.5 U/mL Superase-In [Ambion]), centrifuging at 1000 x g for 5 min at 4 °C after each wash. Cells were then resuspended in Sonication Buffer at a concentration of 100 million per mL and sonicated using a Bioruptor (Diagenode) for 60 cycles (1 cycle: 30 s on, 30 s off) in a 4 °C water bath.

Sonicated extracts were cleared by centrifugation at 16,100 x g for 20 min at 20 °C, then diluted with 1 mL of 1X PAB (8M urea, 100mM HEPES, 200 mM NaCl, 2% sodium dodecyl sulfate) and 3 mL of 2x Hybridization Buffer (1.5 M NaCl, 1.12 M urea, 10X Denhardt's solution, 10 mM EDTA). 200 µL of Dynabeads M-280 Strepavidin magnetic beads (Invitrogen) were rinsed two times with 600 µL ddH$_2$O, then once with 1:2 diluted PAB, added to the extract, and rotated for 1 h at room temperature. Beads were captured twice on a magnetic stand before addition of 4000 pmoles (40 µL) of oligo mixture to the extract (see Supplementary Table S2 for probes used), and overnight hybridization with rotation at room temperature. Oligos for CHART were ordered from IDT and 3′-biotin-TEG, iSp18 spacer modified. The next day, the extract/oligo mixture was centrifuged at 16,100 x g for 20 min at 20 °C to remove insoluble material. 3.2 mL Dynabeads MyOne Streptavidin C1 magnetic beads (Invitrogen) were washed once with 4mL ddH$_2$O and twice with 2 mL 1:2 diluted 1X PAB using the magnetic stand to capture the beads between rinses. Beads were resuspended in 800 µL 1:2 diluted 1X PAB, added to the extract/oligo mixture, and rotated overnight at room temperature. The next day, the beads were captured on a magnetic stand, and 12 mL of Wash Buffer (250 mM NaCl, 10 mM HEPES pH 7.5, 2 mM EDTA, 2 mM EGTA, 0.2% SDS, 0.1% N-lauroylsarcosine) was added, followed by rotation for 2 min. Beads were captured, and 12 mL of Wash Buffer was added again with complete resuspension of beads. This process was repeated an additional 3 times, changing tubes for the final wash. Beads were then washed with 5 mL of Elution Buffer Without Biotin (7.5 mM HEPES pH 7.5, 75 mM NaCl, 1.5mM EDTA, 0.075% N-lauroylsarcosine, 0.15% sodium dodecyl sulfate, 0.02% sodium deoxycholate), then resuspended in 3.2 mL Biotin Elution Buffer (12.5 mM biotin [Invitrogen], 7.5 mM HEPES pH 7.5, 75 mM NaCl, 1.5 mM EDTA,

0.075% N-lauroylsarcosine, 0.15% sodium dodecyl sulfate, 0.02% sodium deoxycholate), followed by 20 min of rotation at room temperature, and a 10 min incubation at 65 °C. The 65 °C incubation was performed using a water bath and the tube was mixed by inversion every 2 min. The supernatant was saved, beads were captured on a magnetic stand, and the entire elution process was repeated once more. Eluents were pooled and 5 mL was added to a Protein Concentrator PES 10K MWCO (Pierce), and spun at 2750 x g at room temperature for 5 min. The final ~1.4 mL was added and the column was spun for an additional 10 min at 2750 x g at room temperature. The concentrated eluent (~120 µL) was removed, and the membrane was washed twice with 100 µL of Biotin Elution Buffer. In our first replicate of CHART mass spectrometry on *Xist*-2kb ESCs, we added 20 µL of RNase H (NEB) to each elution step. For the second replicate of *Xist*-2kb and for the ΔrA234 sample, no RNase H was added in the elution step.

To the concentrated eluents, 25% volume of ice-cold 100% trichloroacetic acid was added, and the sample was incubated overnight at 4 °C with rotation. The sample was then centrifuged at 16,100 x g for 30 min at 4 °C, washed twice with 1 mL of ice-cold acetone with 15 min, 16,100 x g spins at 4 °C after each wash. After the final acetone wash, the pellet was then air-dried and resuspended in 30 µL of NuPAGE 1X LDS Sample Buffer with 5% 2-mercaptoethanol. The sample was boiled for 30 min at 95 °C to reverse crosslinks and loaded onto a NuPAGE 4-12% Bis Tris protein gel (Invitrogen) alongside a Precision Plus Protein Kaleidoscope ladder (Bio-Rad). The gel was then run for 2 cm, rinsed in 50% methanol/10% acetic acid, incubated in fresh 50% methanol/10% acetic acid for 30 min twice, then stained with 0.1% Coomassie Brilliant Blue G-250 (Sigma) in 50% methanol and 10% acetic acid for 3 h. The gel was destained overnight in 50% methanol and 10% acetic acid, and once more with fresh 50% methanol and 10% acetic acid the next day for 3 h. The 2-cm band (invisible by staining) was then excised with a clean razor blade and transferred to a 1.7 mL Eppendorf tube along with a small amount of water, then shipped to the Mass Spectrometry Facility at the University of Massachusetts Medical School for protein identification.

All MS/MS samples were analyzed using Mascot (Matrix Science, London, UK; version Mascot in Proteome Discoverer 2.1.1.21). Mascot was set up to search Uniprot_Mouse assuming the digestion enzyme stricttrypsin. Mascot was searched with a fragment ion mass tolerance of 0.050 Da and a parent ion tolerance of 10.0 PPM. Carbamidomethyl of cysteine was specified in Mascot as a fixed modification. Gln->pyro-Glu of the N-terminus, oxidation of methionine and acetyl of the N-terminus were specified in Mascot as variable modifications. Scaffold (version Scaffold_4.8.0, Proteome Software Inc., Portland, OR) was used to validate MS/MS-based peptide and protein identifications. Peptide identifications were accepted if they could be established at greater than 95.0% probability by the Peptide Prophet algorithm (11) with Scaffold delta-mass correction. Protein identifications were accepted if they could be established at greater than 99.0% probability and contained at least 1 identified peptide. Protein probabilities were assigned by the Protein Prophet algorithm (12). Proteins that contained similar peptides and could not be differentiated based on MS/MS analysis alone were grouped to satisfy the principles of parsimony. Proteins sharing significant peptide evidence were grouped into clusters.

Data for the three samples were compiled in Scaffold and are included as Supplementary File 1. Results were filtered using Scaffold Viewer (4.8.4) with default settings except for the options "Min # Peptides" (changed to 1) and "View→Show Lower Scoring Matches" (toggled off), and Total Spectrum Counts were exported (Supplementary Table S4). Results were further filtered to include proteins which were only present in both *Xist*-2kb replicates, and Total Spectrum Counts were exported (Supplementary Table S3).

***RNA immunoprecipitation***

RNA immunoprecipitation experiments were performed using a modified version of a protocol from (13). ESCs were trypsinized and washed once with PBS before being fixed in 0.3% methanol-free formaldehyde (Pierce) for 30 min with rotation at 4 °C. Formaldehyde was quenched with 125 mM glycine for 5 min at room temperature. ESCs were then washed three times with PBS, snap frozen in liquid nitrogen, and stored at -80 °C.

24 h prior to IP, 5 µL of SPEN (Novus NBP1-82952), RBM15 (Proteintech 10587-1-AP), or 5 µg IgG control (Cell Signaling #3900) antibodies were preconjugated with 25 µL of Protein A/G agarose beads (Santa Cruz) for each IP in a solution of PBS and 0.5% BSA. On the day of the IP, beads were washed once with PBS and 0.5% BSA and once with fRIP Buffer (25mM Tris-HCl pH 7.5, 5mM EDTA, 0.5% IGEPAL CA-630, 150 mM KCl) before being added to sonicated cell lysates.

Prior to sonication, 10 million crosslinked ESCs were resuspended in 0.5 mL RIPA Buffer (50 mM Tris-HCl pH 8, 1% Triton X-100, 0.5% sodium deoxycholate, 0.1% SDS, 5 mM EDTA, 150 mM KCl) containing 0.5 mM DTT, 1:100 protease inhibitor cocktail (Sigma), and 2.5 µL RNasin (Promega) before sonicating using a Vibracell VX130 (Sonics) with two cycles of 30% intensity for 30 s with 1 min of rest on ice between cycles, followed by centrifugation at 4 °C for 15 min at 16,100 x g.

Sonicated ESC lysates were diluted with 0.5 mL fRIP Buffer containing 0.5 mM DTT, 1:100 protease inhibitor cocktail (Sigma), and 2.5 µL RNasin (Promega). A small portion (25 µL) of each lysate was stored at -20 °C for later processing as input. 5 million ESC equivalents were then incubated overnight at 4 °C on a rotating platform with washed, antibody-conjugated beads. The following day, beads were washed once with ice cold fRIP Buffer and then transferred to a clean tube. Beads were then washed 3 times in ice cold ChIP Buffer (50 mM Tris-HCl pH 7.5, 140 mM NaCl, 1 mM EDTA, 1 mM EGTA, 1% Triton X-100, 0.1% sodium deoxycholate, 0.1% SDS), once in ice-cold High Salt Buffer (ChIP Buffer but with 500 mM NaCl) and once in ice-cold LiCl Wash Buffer (20 mM Tris pH 8.0, 1 mM EDTA, 250 mM LiCl, 0.5% NP-40, 0.5% sodium deoxycholate), changing tubes again at the final wash. Each wash (except the first fRIP Buffer wash) was performed for 5 min with rotation at 4 °C. After the final wash, beads were resuspended in ~100 µL 1X Reverse Crosslinking Buffer (1x PBS, 2% N-lauroylsarcosine,

10 mM EDTA) containing 5 mM DTT, 20 µL proteinase K (10 mg/mL Invitrogen 25530015, dissolved in 10 mM Tris pH 7.5, 20 mM $CaCl_2$, 50% glycerol), and 1 µL RNasin (Promega). In parallel, thawed input samples were brought to ~100 µL in the same buffer composition. All tubes were incubated 1 h at 42 °C, 1 h at 55 °C, and 30 min at 65 °C. 1 mL of Trizol was used to extract RNA, and the aqueous phase was supplemented with 1 volume of ethanol and purified using a Zymo-Spin IC column, including on-column DNase I digestion, per the manufacturer's instructions. RNA was eluted in 15 µL dd$H_2O$, and 2 µL from each sample was reverse-transcribed using the MultiScribe High Capacity Kit (Applied Biosystems) with random primers. qPCR was performed using iTaq Universal SYBR Green (Bio-Rad) and custom primers (Supplementary Table S2).

***RBM15/RBM15B/EZH2 knockdown***

Knockdowns of RBM15/RBM15B/EZH2 were generated using two piggyBac-based vectors, one that expresses doxycycline-inducible Cas9 and another used to express sgRNAs targeting exons and the reverse-tetracycline-TransActivator (*rtTA*), modified from (14). Our rationale and construction of these piggyBac-based vectors is described in (15). To create the doxycycline-inducible Cas9 used to cut the DNA of genes of interest, a parent vector was created in which a BGH-poly(A) signal and an EF1α promoter driving expression of a hygromycin resistance gene was ligated into the cumate-inducible piggyBac transposon vector from System Biosciences after its digestion with HpaI and SpeI, which cut just downstream of each chicken β-globin insulator sequence and removed all other internal components of the original vector. The TRE promoter from pTRE-Tight (Clontech) was then cloned upstream of the BGH-poly(A) site, and Cas9 from pX330 (7) was then cloned behind the TRE promoter by digestion with AgeI and SalI (NEB) followed by Gibson Assembly (NEB), to generate the piggyBac cargo vector capable of inducibly expressing Cas9 upon addition of doxycycline. In parallel, individual sgRNAs targeting exons of interest were designed using Desktop Genetics. These sgRNAs were each cloned into BsmBI sites in a U6-sgRNA expression cassette that was inserted upstream of the *rtTA*3-IRES-Neo cassette in the *rtTA*-piggyBac-Cargo vector described in (14). Four sgRNAs per gene of interest were cloned in this manner. For each gene of interest, bacterial cultures expressing each of the four sgRNAs were pooled and grown overnight prior to extraction with PureLink HiPure MidiPrep kit (Invitrogen) of the pooled sgRNA-*rtTA*-expressing plasmids. Sequences of oligos used to create sgRNA-*rtTA*-expressing plasmids are listed in Supplementary Table S2. An sgRNA-*rtTA*-expressing plasmid with an sgRNA insert lacking a targeting sequence was used as a negative control in Figure 4E and F.

To generate TETRISv1 lines with RBM15/RBM15B/EZH2 knockdown, transfections were performed using Lipofectamine 3000 as described above. A total of 2.5 µg of plasmid DNA was transfected per well of a 6-well plate at an 8:2:2:1 ratio of sgRNA-*rtTA* pool : piggyBac-Cas9 : pTETRISv1-*Xist*-2kb : pUC19-piggyBac transposase. Cells were selected for 10 d with 2 µg/mL puromycin, 200 µg/mL G418, and 150 µg/mL hygromycin, starting 24 h after transfection. To induce Cas9 expression and protein knockdown before performing TETRIS assays, cells were treated with 1 µg/mL doxycycline for 4 d. Following this, cells from each TETRIS line were split to 2 wells of a 6-well plate, one with 1 µg/mL doxycycline and one without (for protein lysates for western blot) and to 4 wells of a 24-well plate, two with 1 µg/mL doxycycline and two without (for lysis for luciferase assay).

***Western blots***

For western blot, cells were washed with 1X PBS, harvested by scraping, and lysed by rotating 15 min at 4 °C in RIPA Buffer (10 mM Tris-HCl pH 7.5, 1 mM EDTA, 0.5 mM EGTA, 1% NP-40, 0.1% sodium deoxycholate, 0.1% SDS, 140 mM NaCl) supplemented with 1 mM PMSF and 1:100 protease inhibitor cocktail (Sigma). Lysates were sonicated on ice with a Vibracell VX130 (Sonics) for 2 cycles of 10 sec at 30% output and cleared by centrifuging for 15 min at 16,100 x g at 4 °C before measuring protein concentration using the DC protein assay (Bio-Rad). Equal amounts of protein were run at 150-200 V on 12% polyacrylamide gels with 5% stacking gels and transferred to PVDF membranes at 20 V overnight and at 4 °C. Membranes were blocked with 5% blotting-grade blocker (Bio-Rad) in TBST for 1 h at room temperature. Membranes were then cut to simultaneously probe for a protein of interest (EZH2, RBM15, RBM15B, or luciferase) and a loading control (TBP). Primary incubations were done for 1.5 h at room temperature. Membranes were washed 3 times for 6 min each with TBST at room temperature and incubated with the appropriate secondary antibody for 40 min at room temperature, 1 h at 4 °C, and 20 min at room temperature. Membranes were washed 3 times with TBST for 6 min each and once with TBS for 5 min at room temperature before developing for 5 min at room temperature with Clarity Western ECL Substrate (Bio-Rad; EZH2, luciferase) or SuperSignal West Femto Maximum Sensitivity Substrate (Thermo Scientific; RBM15, RBM15B, TBP) and imaging on a ChemiDoc MP Imaging System (Bio-Rad). Antibodies used: TBP (Abcam ab818; 1:2000 dilution), EZH2 (Cell Signaling Technology CS-5246; 1:1000 dilution), RBM15 (Proteintech 10587-1-AP; 1:1000 dilution), RBM15B (Proteintech 22249-1-AP; 1:1000 dilution), luciferase (Promega G7451; 1:1000 dilution), donkey anti-mouse IgG-HRP secondary (Santa Cruz sc-2314; 1:2500), donkey anti-rabbit IgG-HRP secondary (Santa Cruz sc-2313; 1:2500), donkey anti-goat IgG-HRP secondary (Santa Cruz sc-2020; 1:5000).

***Generation of the* Rosa26 *recombinase-mediated cassette exchange (RMCE) locus by homologous recombination***

A standard *Rosa26* targeting vector was modified to make compatible for recombinase-mediated cassette exchange (RMCE) by creating pR26-RMCE. Briefly, p*Rosa26*-pA was digested with PacI and AscI to insert gene synthesis product SA-Stop-BGHpA-lox2272-mPGK-*PuroRΔTK*-lox66 (BioBasic Inc.). The gene synthesis product SA-Stop-BGHpA-lox2272-mPGK-*PuroRΔTK*-lox66 was released from the commercial vector by KpnI digest and was synthesized to contain 18-bp homologous ends for Gibson assembly into p*Rosa26*-pA. Assembly was performed by mixing equimolar amounts of vector plus insert essentially as described above.

The final targeting vector pR26-RMCE was sequence-verified, linearized by KpnI, and 2.5 μL of 9,750 ng/μL plasmid was nucleofected into approximately 1 million cells of a male F1-hybrid mouse ESC line (derived from a cross between C57BL/6J (B6) and CAST/EiJ (Cast) mice; kind gift of T. Magnuson) using program CG-104 and the Amaxa 4D-nucleofector (V4XP-3024, Lonza). Prior to nucleofection, the ESC line was sent for karyotyping (Karyologic), which verified the 40N, XY nature of the cells. Nucleofected ESCs were plated on three 10-cm plates containing gelatin and gamma-irradiated DR4 mouse embryonic fibroblasts (ASF-1015, Applied Stem Cell). Selection for a successful gene targeting event was performed 48 h after nucleofection with 1 μg/mL puromycin. Long-range PCR was used to screen for homologous recombination across the 3′ *Rosa26* homology arm using 2.5 μL mESC lysate and the thermocycling program (95 °C for 5 min, [98 °C for 20 s, 63 °C for 20 s, 72 °C for 3 min; 40 cycles] 72 °C for 3 min, and 10 °C soak) with primers 6238+ and 6250- (Supplementary Table S2). Two 96-well plates were picked and 10 out of 188 colonies (~5.3%) were positive by long-range PCR.

Genomic DNA was prepared from positive colonies as described in (16) with minor modifications. Briefly, colony DNA was enzymatically digested with 0.5 mg/mL proteinase K in a tail lysis buffer containing SDS. A saturated solution of sodium chloride was added to a final concentration of 25% v/v to salt out protein and centrifuged for 15 min at 10,000 x g. The genomic DNA remained in the supernatant and was transferred to a new tube for isopropanol precipitation, then washed in 70% ethanol to desalt. The DNA sample was resuspended in 150 μL TE with low EDTA (10mM Tris, 0.1mM EDTA, pH 8.0) and allowed to solubilize overnight. 8-10 μg of genomic DNA was digested at 37 °C overnight with 50 U of MscI (R0534L, New England Biolabs). To ensure complete digestion of the genomic DNA, a pilot gel containing 0.7% agarose and TAE was run at 50 V/cm to visualize MscI-cut genomic DNA that appeared as a smear with high-molecular banding. In cases where digestion was incomplete, the DNA was resuspended in 3 volumes of Buffer PB (Qiagen 19066) purified over a silica column, washed with WS Buffer twice (Epoch Life Science B404-400) and eluted in TE with low EDTA (10mM Tris, 0.1mM EDTA, pH 8.0). The incompletely digested DNA was incubated at 37 °C overnight with 2-3 μL containing 10-15 U of MscI in a 50μL reaction to completion. A 15 cm x 10 cm TAE Southern gel with 0.7% agarose was cast and electrophoresed overnight at 20 V/cm to achieve high resolution and efficient separation. The Southern gel was processed as follows: depurination in 0.1 M HCl for 10 min, denaturation twice in 0.1 M NaOH for 15 min, neutralization twice with 100 mM Tris, 100 mM NaCl pH 7.5 for 15 min, and transferred to a wicking apparatus as described by (17) using 20 × SSC to a 0.45 μm microporous nylon 66 membrane on a polyester support, carrying positively charged quaternary ammonium groups (Roche 11417240001). After approximately of 16-24 h of transfer, the wicking apparatus was deconstructed and the transferred DNA was crosslinked to the nylon membrane with 0.12 J of UV light for 90 s. The nylon membrane was blocked by an initial prehybridization step with 30-50 mL of DIG Easy Hyb Granules (Roche 1179689001) for 30 min according to the manufacturer's recommendations. DIG-UTP labeled probes (synthesis below) were denatured for 5 min at 70 °C and then immediately applied to pre-blocked nylon membrane in a rotisserie oven, and hybridization continued overnight with constant rotation. A low and high-stringency wash were each performed at 65 °C (2X SSC, 0.1% SDS for low-stringency; then 0.5 × SSC, 0.1% SDS for high-stringency) for 30 min. Membrane blocking and immunological detection used the DIG Wash and Block Buffer Set (Roche 11585762001) per the manufacturer's recommendations with 1:10,000 dilution of alkaline phosphatase-conjugated anti-DIG antibody and with 1:100 dilution of chemiluminescent substrate CDP-Star (Anti-digoxigenin-AP conjugate, Fab frag, Roche 11093274910; CDP-Star, Roche 11759051001).

Southern blotting probes were labeled by PCR with a dNTP mix containing 200 μM dATP, dGTP, dCTP; 130 μM dTTP and 14-70 μM alkali-labile digoxigenin-dUTP according to the manufacturer's recommendations (PCR DIG Probe Synthesis Kit, Roche 11636090910). Primers for probe synthesis are described in Supplementary Table S2.

#### *Construction of pCARGO-RMCE, pCARGO-RMCE containing the complete* **Xist** *genomic locus, pCARGO-RMCE-***Xist***-5.5kb, pCARGO-RMCE-***Xist***-2kb+6,7, and pCARGO-RMCE-***Xist***-2kb*

The base vector pCARGO-RMCE was constructed from pLCA.66/2272 and gene synthesis product lox71-TRE-CMV-mcs-SV40pA-FRT-mPGK-Em7-Hygro-FRT-lox2272 (BioBasic Inc,.). The vector pLCA.66/2272 was a gift from M. Magnuson (Addgene plasmid #22733; (18)) and digested with AatII and XhoI. The insert lox71-TRE-CMV-mcs-SV40pA-FRT-mPGK-Em7-Hygro-FRT-lox2272 was digested with AatII and XhoI and had 18-bp compatible ends to vector pLCA.66/2272, and the insert cassette was subcloned by Gibson assembly. Assembly was performed by mixing equimolar amounts, e.g. 0.125 pmol each, of vector plus insert in a final volume of 20 μL with 2x assembly mix and incubating at 37 °C for 7:30, 50 °C for 15 min, 50 °C for 1 min where -1 °C/cycle, n=10 cycles, 50 °C for 35 min, and final soak at 10 °C. To make 2x assembly master mix as modified from (19), the following reagents were prepared as follows: 6 mL of 5x isothermal buffer (3 mL 1 M Tris, pH 7.5, 150 μL 2 M $MgCl_2$, 600 μL of 40 mM lithium salt-dNTP mix, 300 μL of 1 M TCEP pH 7.5, 1.5 g of PEG-8000, 300 μL of 100 mM NAD, and Milli-Q water to 6 mL). The final 2X assembly mix was prepared to a volume of 600 μL by addition of 240 μL of 5X isothermal buffer, 2.4 μL of 10 U/μL T5 FEN, 15 μL of 2 U/μL PfuX7, 120 μL of 40 U/μL Taq DNA ligase, and Milli-Q water to 600 μL (19).

The pCARGO-RMCE-*Xist*-RFP capture vector was generated from pCARGO-RMCE by double digestion with MluI and SwaI and inserting PCR generated amplicons for *Xist* homology arms, BBa_292001, ccdB, and RFP into the vector by Gibson assembly using equimolar DNA parts of 0.125 pmol each and transforming into Survival2 cells (A10460, Thermo Fisher). The *Xist* 5′ homology arm was 162 bp, and the 3′ homology arm was 416 bp. Sequence verification of multiple clones could not identify any clones with a functional ccdB, and the final vector contained a frameshifting ccdB deletion. The red fluorescent protein mScarlet-I was functional and driven by the double terminator + constitutive promoter J23100 from BBa_292001. RFP was used for visual inspection of background transformants, e.g. red colonies, after recombineering. Plasmid DNA purified by the standard miniprep method of alkaline lysis with NaOH as described originally by Birnboim and Doly, contains approximately 3% cyclic coiled DNA that is resistant to restriction digestion and leads to background transformants (20). The final capture vector pCARGO-RMCE-*Xist*-RFP was digested with PacI and gel extracted for retrieval of lncRNA *Xist* by gap repair.

An *Xist* fosmid from the WIBR-1 mouse library (*Mus musculus*, female, strain C57BL/6J) corresponding to clone number WI1-2121K18 was obtained through BACPAC Resources. The original fosmid library was constructed by shearing genomic DNA and ligating into the Eco72I-linearized pEpiFOS-5 vector and transformed into the host *E. coli* strain Epi100. To make Epi100 proficient for recombineering, the *Xist* fosmid clone was infected with a replication-defective λ phage containing *exo*, *bet* and *gam* under the control of its native phage operon containing the pL promoter and temperature-sensitive repressor, cI$_{857}$ (21,22). Briefly, Epi100 cells containing the *Xist* fosmid were grown to saturation in LB-Lennox broth containing 1% maltose and 12.5 µg/mL chloramphenicol, and diluted the following morning 70-fold until OD$_{600}$ = 0.1. 3 mL of culture was grown to the exponential phase OD$_{600}$ = 0.6, pelleted by centrifugation, then washed with 1 mL 10 mM $MgSO_4$, resuspended in 100 µL 10 mM $MgSO_4$ with 10 µL replication-defective λ phage (λ cI$_{857}$ ind1 Cro$_{TYR26amber}$ P$_{GLN59amber}$ rex< >tetRA), and then incubated with shaking at 32 °C for 1 h. The lysogenization frequency is typically 1%; therefore, selection for clones containing both the fosmid and stable integration of the defective prophage is done by plating dilutions of clones on 2xYT agar plates containing 12.5 µg/mL chloramphenicol and 10 µg/mL tetracycline at 32 °C. Clones containing the stable-integrated lysogen will be tetracycline resistant and do not express the prophage exo, bet and gam at 32 °C, but can be induced to make the recombineering proteins conditionally by shifting the temperature of bacteria to 42 °C. For production of a stock of λ phage lysate, bacterial strain LE392 (K9981, Promega) containing lysogen (λ cI$_{857}$ ind1 Cro$_{TYR26amber}$ P$_{GLN59amber}$ rex< >tetRA) was induced to produce phage by shifting an exponentially growing culture in terrific broth to 42 °C for 15 min, and lysing by then shifting to 39 °C for 1 h containing 50 µL BCP (B9673-200ML, Millipore Sigma) (21).

The final pCARGO-RMCE-*Xist* was generated by transforming 3 µL of 300 ng linearized capture vector pCARGO-RMCE-*Xist*-RFP into recombineering-proficient WI1-2121K18 after 42 °C induction for 15 min and washes to make cells electrocompetent essentially as described by (22). Recombineered clones were visually inspected for only white colonies, then screened for a 5′ recombinant and 3′ recombinant junction generated by a successful gap repair event (data not shown), and 33 of 66 clones were positive by PCR screening. Ten clones were miniprepped and digested with PstI-HF or triple cut with PacI/AsiSI/AscI, and 2 out of 10 clones were correct by restriction mapping. Correctly retrieved clones also contained the parental BAC as evidenced by streaking on 12.5 µg/mL chloramphenicol 2xYT plates. Plasmid DNA was purified by standard silica-column purification and 1 ng of pCARGO-RMCE-*Xist* was transformed into 40 µL electrocompetent Epi300 (EC300110, Lucigen). A representative clone was restreaked on a 12.5 µg/mL chloramphenicol 2xYT plate and was antibiotic sensitive. This clone was used to prepare transfection quality plasmid DNA for RMCE using the NucleoBond BAC 100 purification kit (Macherey-Nagel).

Independently, the first 2016 nucleotides of *Xist* (*Xist*-2kb) was PCR amplified and cloned into the MluI-HindIII sites in pCARGO-RMCE by Genewiz, Inc. Similarly, exon6-intron-exon7 was PCR amplified from the *Xist* fosmid (WI1-2121K18) and cloned into the AatII and XhoI sites in pCARGO-RMCE-*Xist*-2kb by Genewiz. The first 5588 nucleotides of *Xist* (*Xist*-5.5kb) was PCR amplified from the *Xist* fosmid (WI1-2121K18) using (NEBNext HF 2X master mix, NEB M0541S) and cloned into the ClaI sites in pCARGO-RMCE. Note that not all of 5588 nucleotides of the endogenous *Xist* sequence were successfully cloned into the vector – see description in results section as well as Figure 5A and Supplementary Figure S6A. The pCARGO-*Xist*-2kb, pCARGO-*Xist*-5.5kb, and pCARGO-*Xist*-2kb+6,7 plasmids were prepared for electroporation using the PureLink HiPure MidiPrep kit (Invitrogen).

***Generation of clonal ESCs inducibly expressing* Xist*-2kb,* Xist*-5.5kb,* Xist*-2kb+6,7, and full-length* Xist *from the* Rosa26 *locus***

To insert sequences carried in pCARGO into the *Rosa26* locus of the RMCE cell line described above, the pCARGO plasmid (empty pCARGO [6.25 µg], pCARGO carrying *Xist*-2kb [6.25 µg], or full-length *Xist* [19 µg]) was coprecipitated with pOG-Cre at an approximate ratio of 1:2.4 (pOG-Cre:pCARGO) by plasmid copy number, along with 6 µg of an additional plasmid containing a hygromycin resistance gene (to provide transient hygromycin resistance). For *Xist*-5.5kb and *Xist*-2kb+6,7, the pCARGO plasmid (pCARGO-*Xist*-5.5kb [9.5 µg] or pCARGO-*Xist*-2kb+6,7 [18.83 µg]) was coprecipitated with pOG-CRE at an approximate ratio of 1:1 by plasmid copy number. The plasmids were precipitated and resuspended in 10 µL TE and mixed with 1 million *Rosa26*-RMCE cells for electroporation using the Neon Transfection System (Invitrogen) with a 100 µL pipette tip. Electroporation conditions were 1 pulse, 40 ms, 1000 V. Following electroporation, cells were plated on a 10-cm plate seeded with 1.5 million gamma-irradiated multi-drug resistant DR4 MEFs (ATCC SCRC-1045) in ESC medium without penicillin/streptomycin. Approximately 48 h after selection, medium was changed to normal ESC medium with penicillin/streptomycin. At 72 h, hygromycin was added to the medium at 150 µg/mL. After 4 d of hygromycin selection, 3 µM ganciclovir was added along with the hygromycin. Colonies were picked when ready (approximately 4 d after adding ganciclovir, or 11 d after electroporation) and maintained on gamma-irradiated DR4 MEFs with hygromycin and ganciclovir.

Freezer stocks and genomic DNA were prepared from each colony. Genomic DNA (gDNA) was prepared by incubating cells overnight at 55 °C in 400 µL ESC lysis buffer (100 mM Tris-HCl, pH 8.0, 5 mM EDTA, pH 8.0, 200 mM NaCl, 0.2% SDS) per well of a 24-well plate supplemented with 80 µL Proteinase K (20 mg/mL; Denville Scientific) and 8 µL linear acrylamide, followed by 1 h at 100 °C. gDNA was precipitated by adding 960 µL cold 100% ethanol, rotating 15 min at 4 °C, and spinning at 16,100 x g for 5 min at 4 °C. The gDNA pellet was washed with 1 mL 80% ethanol and resuspended in 100 µL TE overnight at room temperature.

Genotyping PCR reactions were performed with gDNA using Apex Taq DNA Polymerase (Genesee Scientific). Primer pairs (RMCE_PCR1 and RMCE_PCR2 in Supplementary Table S2) either amplified a product in the non-recombined Rosa26 target locus (no insertion from pCARGO) or in the recombined locus (pCARGO inserted).

To generate RMCE cells inducibly expressing the inserted pCARGO sequences, an *rtTA* gene was inserted using piggyBac-mediated transgenesis. Briefly, 400,000 RMCE cells were seeded in a single well of a 6-well plate and transfected 24 h later with 0.5 µg *rtTA*-cargo plasmid and 1 µg of pUC19-piggyBac transposase plasmid (14) using Lipofectamine 3000 (Invitrogen) according to manufacturer instructions. Cells were selected for 7-9 days with G418 (200 µg/mL) beginning 24 h after transfection. Inducible pCARGO sequence expression was verified by RT-qPCR and RNA FISH.

#### ***RNA sequencing and analysis***

Replicates for RNA-seq were as follows: for *Rosa26*-RMCE empty-cargo and *Xist*-2kb cells, fractionated RNA was prepared from 2 independent clones, one independently induced with doxycycline twice and one induced once (3 replicates each); for *Rosa26*-RMCE *Xist*-5.5kb and *Xist*-2kb+6,7 cells, fractionated RNA was prepared from 2 independent clones each induced once (2 replicates each), and for *Xist*-full length cells, fractionated RNA was prepared from 1 clone independently induced twice (2 replicates).

RNA-seq libraries were prepared using the RNA HyperPrep Kit with RiboErase (Kapa Biosciences) and sequenced on an Illumina NextSeq 500 machine using a 75-cycle high output NextSeq kit (Illumina). Sequencing reads were aligned to mm9 genomic sequence using Star (version 2.5.4b; (23)) with default parameters. All mm9 genome annotations were obtained from the UCSC genome browser (24). Variant sequence data were obtained from the Sanger Institute (http://www.sanger.ac.uk/resources/mouse/genomes/). Only reads that had a mapping quality greater than or equal to 30 were used. CAST/EiJ pseudogenome creation and allele-specific read retention was performed as in (25,26). Reads were assigned to genes using featureCounts (27). Differential expression analysis (comparing allelic cytoplasmic counts between empty pCARGO and *Xist* transgene lines) was done using DESeq2 (28). MA plots were generated using the plotMA function in DESeq2 with default settings after shrinking log2fold changes (lfcShrink function in DESeq2). To quantify nuclear fraction of transcripts, upper quartile-normalized nuclear and cytoplasmic counts for each gene were added together and the normalized nuclear count values were divided by the total. All genome-related plots were generated using R (version 3.4.4; (29)).

***Visualization of predicted and observed polyadenylation sites within*** **Xist**

*Xist* genomic sequence (RefSeq NR_001463.3, 213742) plus the 1000 bp immediately downstream was retrieved from the UCSC Genome Browser and used to query PolyA_SVM (version 2.2; http://exon.umdnj.edu/polya_svm_server/index.html) for predicted polyadenylation sites, using default settings except with mouse as the training species model. PolyA_SVM sites were converted to chromosomal coordinates (mm9 chrX), which were used to generate a BED file. PolyA_SVM scores (range 0-6) were converted to BED scores (range 0-1000) so that higher BED scores correspond to lower (stronger) PolyA_SVM scores: BED_score = 1000 - 1000*(PolyA_SVM score)/6.

Experimental polyadenylation sites for mouse *Xist* (RefSeq 213742) were retrieved from PolyA_DB (version 3.2; http://exon.umdnj.edu/polya_db/v3/). A BED file was created using the PolyA_DB chromosomal coordinates and BED score calculated from the PolyA_DB Mean RPM value (range 0.4-36.7 within *Xist*) as follows: BED_score = 1000*((Mean_RPM)-0.4)/36.7.

In the UCSC Genome Browser (mouse mm9), custom tracks were generated using the PolyA_SVM and PolyA_DB BED files, displayed alongside built-in PolyA-Seq tracks with default settings.
